# On the relationships between rarity, uniqueness, distinctiveness, originality and functional/phylogenetic diversity

**DOI:** 10.1101/2021.08.09.455640

**Authors:** Sandrine Pavoine, Carlo Ricotta

## Abstract

Rarity reflects the low abundance of a species while distinctiveness reflects its quality of being easy to recognize because it has unique functional characteristics and/or an isolated phylogenetic position. As such, the assemblage-level rarity of a species’ functional and phylogenetic characteristics (that we name ‘effective originality’) results from both the rarity and the distinctiveness of this species. The functional and phylogenetic diversity of an assemblage then results from a compromise between the abundances and the effective originalities of the species it contains. Although the distinctiveness of a species itself depends on the abundance of the other species in the assemblage, distinctiveness indices that are available in the ecological literature scarcely consider abundance data. We develop a unifying framework that demonstrates the direct connections between measures of diversity, rarity, distinctiveness and effective originality. While developing our framework, we discovered a family of distinctiveness indices that permit a full control of the influence one wants to give to the strict uniqueness of a species (=its smallest functional or phylogenetic distance to another species in the assemblage). Illustrating our framework with bat phylogenetic diversity along a disturbance gradient in Mexico, we show how each component of rarity, distinctiveness and originality can be controlled to obtain efficient indicators for conservation. Overall our framework is aimed to improve conservation actions directed towards highly diverse areas and/or towards species whose loss would considerably decrease biodiversity by offering flexible quantitative tools where the influence of abundant versus rare, and ordinary versus original, species is understood and controlled.

## 1. Introduction

Generally speaking, biological diversity or biodiversity is the range of many different characteristics of biological systems. In species assemblages, biodiversity thus emerges because species are not equivalent; and species diversity increases with species richness (the number of species) and the evenness (or equitability) of species abundances (e.g., Patil and Taillie, 1982). In local assemblages, often many species are rare, having small population size, and only a few species dominate in abundance (e.g., Hughes, 1986), yielding moderate levels of species diversity. However the exact shape of species abundance distribution may depend on ecological processes as for example disturbance can modify species relative abundances, and consequently species diversity (e.g., Matthews and Whittaker, 2015).

Another increasingly studied aspect of species rarity that may influence biodiversity levels is the rarity of species biological characteristics (Kondratyeva et al., 2019). Pavoine et al. (2017) defined the originality of a species in a given assemblage as ‘the rarity of its biological characteristics’. As such, originality can be based on the species evolutionary history (phylogenetic originality) or on the species functional traits (functional originality) as both aspects are believed to reflect biological differences between species. Originality is related in the ecological literature to the concepts of species distinctiveness and uniqueness. Uniqueness is the quality of being unique in some way (Cambridge Dictionary, 2021), here in the functional characteristics or the phylogenetic position. Distinctiveness is the quality of being easy to recognize because of being different from other things (Cambridge Dictionary, 2021). In our context, the distinctiveness of a species can thus be defined as its quality of being easy to recognize because it has some unique functional characteristics or as its quality of being easily found in a phylogenetic tree because it belongs to an old, species-poor clade. Distinctiveness and uniqueness are thus two closely connected concepts as the more unique a species is the more distinct it might be. To simplify the writing, hereafter we will thus use below only the term distinctiveness. We will also use the acronym ‘FP’ to mean ‘functional or phylogenetic’.

Originality is the quality of being special and interesting and not the same as anything or anyone else (Cambridge Dictionary, 2021). This is why the originality of a species can be considered as the assemblage-level rarity of the FP-characteristics associated to this species (Pavoine et al., 2017). This originality depends on whether abundance data are considered (Fig. 1). In absence of abundance data, the concept of originality is equivalent to that of distinctiveness (Pavoine et al., 2005). This is because the species own rarity is discarded and the rarity of a species’ biological characteristics is only linked to the proportion of functional traits or phylogenetic history that are unshared with other species (see, e.g., the distinct species 1 and 2 in Fig. 1a,b,c). In global conservation studies for example abundance data are rarely considered (e.g., Gaüzère et al., 2015).

**Fig. 1.**
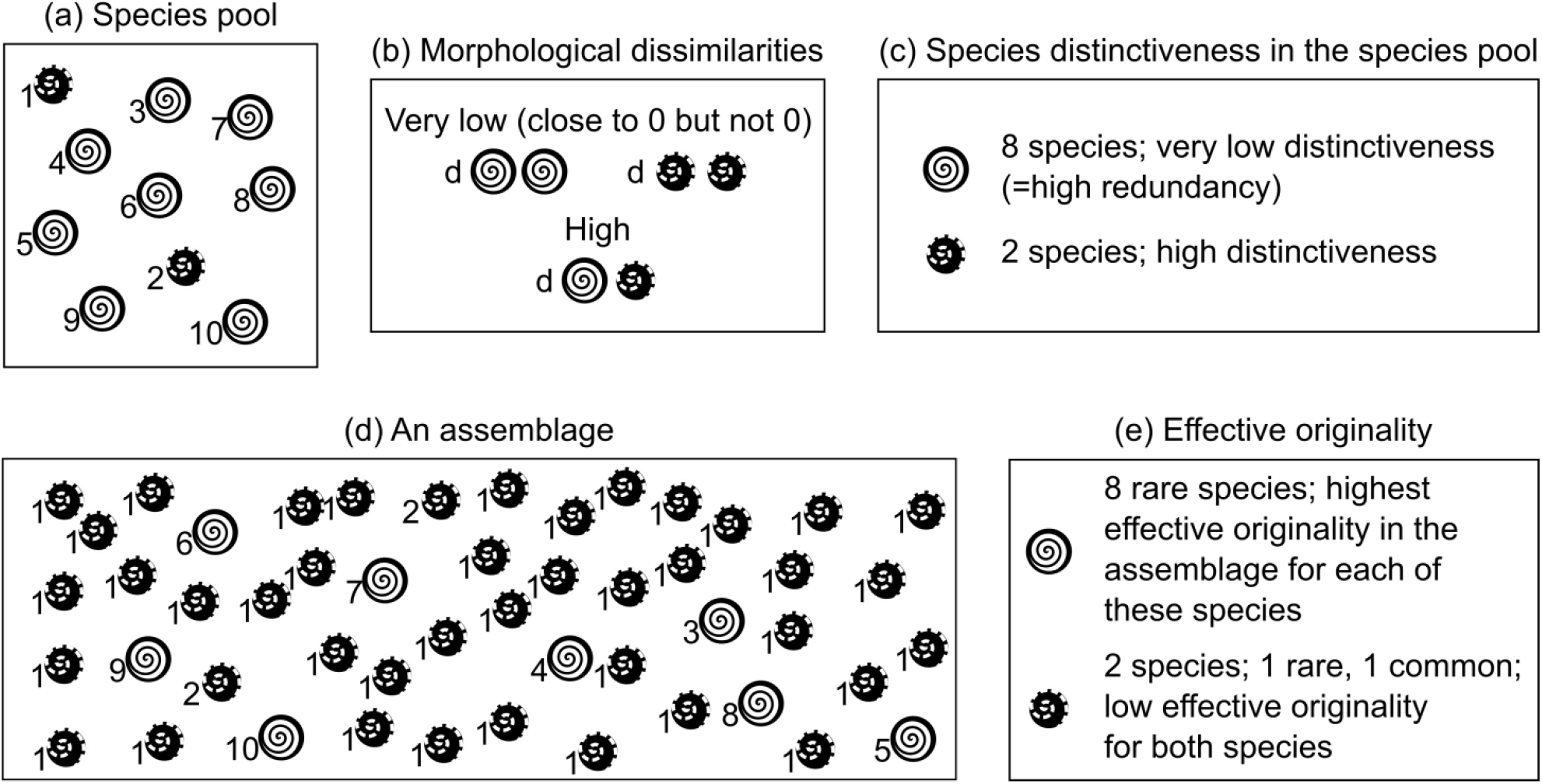
Theoretical illustration of the concept of effective originality. We considered 10 theoretical species that form the species pool (panel a). We considered the morphological aspects of the species, imposing (panel b) that none of the species is exactly similar to another in the pool but that the species represented by the same symbols share many morphological characteristics. Two species are black with white broken lines and have the most distinct morphology in the species pool (panel c). We considered an assemblage where species numbered 1 is 30 times more abundant than all other species (panel d). Due to the skewed distribution of abundance, in this assemblage the white species with black lines have the highest effective originality (panel e).

For more local ecological studies however, abundance data are often available and often reveal meaningful to analyze ecological systems (e.g. Enquist et al., 2019). Consider that a focal species *j* is distinct from other species in a defined species pool. If in an assemblage, its few functionally sibling or close relatives “dominate” the assemblage by their high abundance or if species *j* is itself very abundant, then the assemblage-level rarity of the FP-characteristics associated to species *j* is actually low and the originality of this species is in fact low (e.g., species 1 and 2 in Fig. 1d,e). Inversely, consider that species *j* has low distinctiveness in the species pool (where abundance data is discarded). Species *j* may still be effectively original in an assemblage where its abundance and those of its sibling, close species are all low (e.g. species 3 to 8 in Fig. 1d,e). Hereafter, we will refer to this abundance-based definition of FP-originality as the effective FP-originality. Combining the different aspects of a species rarity (its low abundance; the distinctiveness of its traits; its isolated position in the phylogeny), for the scope of this paper, we will thus consider two intuitive conditions that a measure of effective FP-originality should respect. Consider an assemblage composed of *N* species with relative abundances *p _j_* (*j* = 1, 2,…, *N*), with 0 < *p _j_* ≤ 1 and 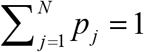. The two conditions are:

C1. the effective originality of a given species *j* should increase with its functional or phylogenetic distinctiveness with respect to the individuals of other species in the assemblage, and
C2. the effective originality of a given species *j* should increase with its abundance-based rarity, a decreasing function of *p _j_*.

In our context, the effective originality of a species can thus be considered as its quality of being of ecological interest because it is special, all both rare, unique and distinct from all other species in an assemblage. An example of effectively original species is the critically-endangered Western Swamp Tortoise (*Pseudemydura umbrina*) endemic to Western Australia (Bouma et al., 2020), morphologically unique among extent Chelidae. The originality of this species may be due to strict trait conservatism along its evolution as fossil data indicates little morphological change for this species since the early Miocene (Zhang et al., 2017). The small population size of this endemic species (Arnall, 2018) could yield a low contribution of this species to Chelidae biodiversity. However, in contrast, due to its distinct morphological characteristics, the theoretical contribution of this species to the biodiversity of the Chelidae family is expected to be high. Moreover, its low abundance reinforces the rarity of its morphological characters and thus the key necessity to consider this species in conservation actions. Indeed, it has already been shown that rare species (of low abundance) may contribute more to ecosystem functioning and ecosystem services than their low abundance would suggest because of their unique functional role, notably via unique functional traits (Dee et al., 2019). While, if species abundance is not considered, the contribution of a species to biodiversity can directly result from its distinctiveness, considering abundance yields its contribution to result from a compromise between its distinctiveness and its abundance. However, although many species, trait and phylogenetic diversity measures do include abundance data, very few originality measures developed so far include species abundance; and the full potential, for ecological studies, of these few abundance-weighted measures of originality still needs to be emphasized.

We thus focus our study on three of these abundance-weighted measures of originality: Cadotte et al. (2010) *AED* index and Kondratyeva et al. (2019) *O* index for phylogenetic originality; and Ricotta et al. (2016) *K* index for functional originality. These measures of originality have the advantage to be linked with standard measures of diversity: Faith phylogenetic diversity (PD; Faith 1992) for the measure of Cadotte et al., an abundance-weighted generalization of Faith’s PD (Pavoine et al., 2009) for the measure of Kondratyeva et al. and Rao’s quadratic entropy (hereafter more simply named quadratic diversity; Rao, 1982) for the measure of Ricotta et al. We extend below these three originality indices and unify them in a common framework on the link between diversity, rarity, distinctiveness and originality.

## 2. Quadratic diversity as a mean of effective originalities

Let *d_ij_* be the FP-dissimilarity between species *i* and *j* such that *d_ij_* ≥ 0 and *d _jj_* = 0. Dealing with FP-diversity, we can define the FP-distinctiveness of species *j* as the weighted mean FP-dissimilarity of *j* from all other species in the assemblage (Ricotta et al., 2016):

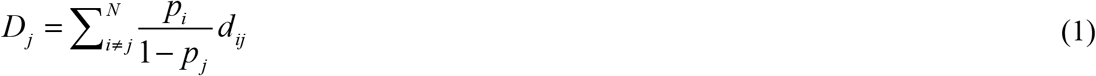

where the quantity *p_i_*/(1 − *p_j_*) is the relative abundance of species *i* (*i* ≠ *j*) with 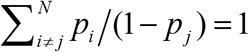. *D _j_* satisfies condition C1 of a measure of effective originality but not condition C2. A community-level measure of expected FP-distinctiveness defined as 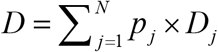 would not behave as an index of diversity (see Appendix A for details).

An index of FP-diversity can however be obtained by combining a species abundance-based rarity with its FP-distinctiveness. For *d_ij_* in the range [0, 1], consider *s_ij_*, the similarity between species *i* and *j* calculated as *s_ij_* = 1 − *d_ij_*. Let 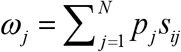 be the ordinariness of species *j*, i.e., the expected similarity between an individual of species *j* and an individual chosen at random in the assemblage, including the individuals of species *j* itself. According to Leinster and Cobbold (2012), *ω_j_* can be thus interpreted as the relative abundance of all species that are FP-similar to *j*. Since 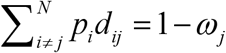, we have:

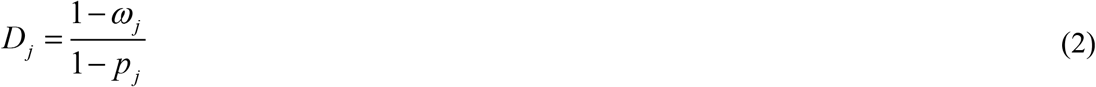

The denominator of Eq. 2, *ρ_j_* = 1 − *p_i_*, is the rarity of species *j* according to the well-known Simpson diversity (Simpson, 1949; Patil and Taillie, 1982). The numerator of Eq. 2, *O_j_* = 1 − *ω _j_* is identical to the species-level originality index *K_j_* of Ricotta et al. (2016) that we mentioned in the introduction. *O_j_* satisfies both conditions C1 and C2 of an index of effective originality. The corresponding community-level measure of expected originality equals Rao’s quadratic diversity:

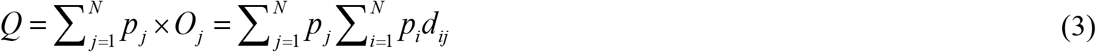

the abundance-weighted average FP-dissimilarity between any two species in the assemblage. In contrast to FP-distinctiveness *D_j_*, effective FP-originality *O_j_* accounts for the abundance of species *j* itself: to evaluate the rarity of the biological characters of a focal species when abundance data are considered, then the abundance of the focal species itself has to be considered.

## 3. Parametric generalizations

We consider below, two possible parametric extensions of Rao’s quadratic diversity. The parameter α of the first one controls the importance given to ordinary species in opposition to effectively original species:

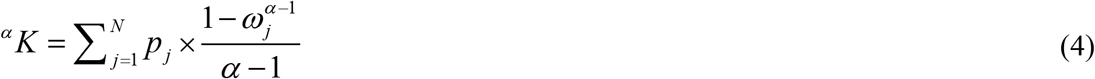

Eq. 4 was first developed by Ricotta and Szeidl (2006). For *α* = 2, ^2^*K* = *Q* and for α tending to 1, it is a generalization of the Shannon index (Shannon, 1948; Ricotta and Szeidl, 2006). When parameter *α* in *^α^K* increases, then ordinary species (those species with low effective originality according to the quadratic diversity) are given increasingly important weights in the measurement of phylogenetic diversity. We develop in Table 1 a decomposition of *^α^K* in terms of rarity, distinctiveness and originality (see details and proofs in Appendix A).

**Table 1.** Summary of the links between rarity, distinctiveness, originality, and diversity developed in the main text with diversity indices *Q*, *^α^K* and *^α^K*\*.

|  |  | Quadratic diversity<sup>a</sup> | First parametric extension<sup>a</sup> | Second parametric extension<sup>a</sup> |
| --- | --- | --- | --- | --- |
| Species-level |  |  |  |  |
| | FP-distinctiveness | $D_j = \sum_{i \neq j}^N \frac{p_i}{1 - p_j} d_{ij}$ | ${}^\alpha D_j = \frac{1 - \left( \sum_{i=1}^N p_i s_{ij} \right)^{\alpha-1}}{1 - p_j^{\alpha-1}}$ | ${}^\alpha D_j^* = \frac{\sum_{c=1}^{N-1} u_{c\vert j} (1 - p_{c\vert j}^{\alpha-1})}{(1 - p_j^{\alpha-1})}$ |
| | abundance-based rarity | $\rho_j = (1 - p_j)$ | ${}^\alpha \rho_j = \frac{1 - p_j^{\alpha-1}}{\alpha - 1}$ | ${}^\alpha \rho_j = \frac{1 - p_j^{\alpha-1}}{\alpha - 1}$ |
| | effective originality | $O_j = U_j \times \rho_j$ | ${}^\alpha O_j = {}^\alpha D_j \times {}^\alpha \rho_j$ | ${}^\alpha O_j^* = {}^\alpha D_j^* \times {}^\alpha \rho_j$ |
| Community-level |  |  |  |  |
| | species diversity<sup>b</sup> | $S = \sum_{j=1}^N p_j \rho_j$<br>$= 1 - \sum_{i=1}^N p_i^2$ | ${}^\alpha S = \sum_{j=1}^N p_j ({}^\alpha \rho_j)$<br>$= \frac{1 - \sum_{j=1}^N p_j^\alpha}{\alpha - 1}$ | ${}^\alpha S = \sum_{j=1}^N p_j ({}^\alpha \rho_j)$<br>$= \frac{1 - \sum_{j=1}^N p_j^\alpha}{\alpha - 1}$ |
| | FP-diversity | $Q = \sum_{j=1}^N p_j O_j$<br>$= \sum_{i=1}^N \sum_{j=1}^N p_i p_j d_{ij}$ | ${}^\alpha K = \sum_{j=1}^N p_j ({}^\alpha O_j)$<br>$= \sum_{j=1}^N p_j \times \frac{1 - \left( \sum_{i=1}^N p_i s_{ij} \right)^{\alpha-1}}{\alpha - 1}$ | ${}^\alpha K^* = \sum_{j=1}^N p_j ({}^\alpha O_j^*)$<br>$= \sum_{j=1}^N p_j \times \frac{\sum_{c=1}^{N-1} u_{c\vert j} (1 - p_{c\vert j}^{\alpha-1})}{\alpha - 1}$ |
| | Species equivalents<sup>c</sup> | $E = 1 / \sum_{i=1}^N \sum_{j=1}^N p_i p_j s_{ij}$ | ${}^\alpha E = \left( \sum_{j=1}^N p_j \left( \sum_{i=1}^N p_i s_{ij} \right)^{\alpha-1} \right)^{\frac{1}{(1-\alpha)}}$ | ${}^\alpha E^* = \left( \sum_{j=1}^N p_j \left( 1 - \frac{\max_i (d_{ij})}{d_{\max}} + \sum_{c=1}^{N-1} \frac{u_{c\vert j}}{d_{\max}} p_{c\vert j}^{\alpha-1} \right) \right)^{\frac{1}{(1-\alpha)}}$ |

<sup>a</sup> $Q = {}^2K^* = {}^2K^*$ ; FP-components are expressed in terms of dissimilarities ( $d_{ij} \geq 0$ ) or similarities ( $s_{ij} = 1 - d_{ij} / d_{\max}$ , with $d_{\max} \geq \max_{ij}(d_{ij})$ ) between species; Notations are identical as in the main text; see Table 2 for the special case of phylogenetic dissimilarities.
<sup>b</sup> $S$ = Simpson index (Simpson, 1949; Patil and Taillie, 1982); ${}^\alpha S$ = HCDT index (see e.g., Kondratyeva et al., 2019).
<sup>c</sup> See Leinster and Cobbold (2012) who first developed a function similar to ${}^\alpha E$ .

We develop here the following alternative to index *^α^K*, named *^α^K*\*, where parameter *α* controls species’ abundance instead of ordinariness, i.e. *α* controls the importance given to abundant species in opposition to rare species. As such, with *^α^K*\*, varying parameter *α* influences the ranking of species according to their distinctiveness and effective originality:

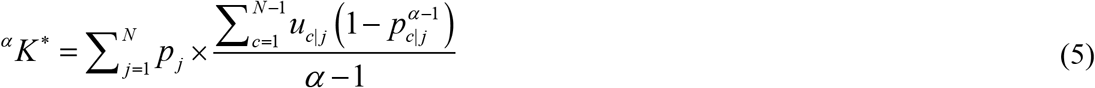

where *c*|*j* indicates the *c*th closest species from species *j*; *u*_*c*|*j*_ = *d*_*c*|*j*,*j*_ − *d*_*c*−1|*j*,*j*_ for *c* > 1 with 0|*j* = *j* and thus *d*_0|*j*,*j*_ = 0 and *p*_0|*j*_ = *p_j_*; and 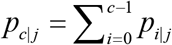. Compared with *^α^K*, *^α^K*\* does not require that the *d_ij_* vary in [0,1]; the only necessary condition is *d_ij_* ≥ 0. For *α* = 2, ^2^*K* * = *Q*. For α tending to 1, it is a generalization of the Shannon index different from that associated with index *^α^K* (Appendix A). We provide in Table 1 a decomposition of *^α^K* in terms of rarity, distinctiveness and originality (see details and proofs in Appendix A).

For the specific case of species characterized by their rooted phylogenetic tree, the parametric index *^α^I* of phylogenetic diversity proposed by Pavoine et al. (2009) (Table 2) is equivalent to *^α^K*\* applied to *d_ij_* = the sum of branch lengths on the shortest path from tip *j* to its most recent common ancestor with species *i*. In addition, in this particular case the associated species effective originalities (Table 2) are equivalent to those introduced by Kondratyeva et al. (2019) who expressed index *^α^I* of phylogenetic diversity as a mean of the species originalities. For *α* = 0, the phylogenetic effective originality associated with *^α^I* (Table 2) reduces to *AED_j_* × *n* − *H _j_* (Kondratyeva et al. 2019) where *AED* is Cadotte et al. (2010) index that we referred to in the introduction, *n* is the total number of individuals in the assemblage and *H _j_* is the sum of branch lengths from the focal species *j* to the root of the tree. This links, in a unified framework, Ricotta et al. *K*, Kondratyeva et al. *O* and Cadotte et al. *AED* measures of abundance-weighted originality. *^α^I* provides thus a consistent alternative to *^α^K*\* in the special case of a phylogenetic tree and we depict its writing in terms of rarity, distinctiveness and effective originality in Table 2. Similarly, we introduce in Table 2 a rewriting (named *^α^Y*) of index *^α^K* and its associated indices of distinctiveness and originality, in the special case where species are tips of a phylogenetic tree and the phylogenetic distance between species *j* and *i* is calculated as the sum of branch lengths on the shortest path from tip *j* to its most recent common ancestor with species *i*. All functions of rarity, distinctiveness and effective originality discussed here are thus connected in a global framework in Tables 1 and 2, highlighting the strong links between different facets of rarity, distinctiveness, originality and diversity.

**Table 2.** Adaptation of Table 1 in the case where species are tips of a phylogenetic tree.

|  | First parametric extension<sup>a</sup> | Second parametric extension<sup>a</sup> |
| --- | --- | --- |
| Species-level |  |  |
| distinctiveness | ${}^{\alpha}\Delta_j = \frac{1 - \left( \sum_{b \in C(j, \text{root})} \frac{L_b}{H} p_b \right)^{\alpha-1}}{1 - p_j^{\alpha-1}}$ | ${}^{\alpha}\Delta_j^* = \frac{\sum_{b \in C(j, \text{root})} L_b (1 - p_b^{\alpha-1})}{1 - p_j^{\alpha-1}}$ |
| effective originality | ${}^{\alpha}\Omega_j = {}^{\alpha}\Delta_j \times {}^{\alpha}\rho_j$ | ${}^{\alpha}\Omega_j^* = {}^{\alpha}\Delta_j^* \times {}^{\alpha}\rho_j$ |
| Community-level |  |  |
| FP-diversity<sup>b</sup> | $\begin{aligned} {}^{\alpha}Y &= \sum_{j=1}^N p_j ({}^{\alpha}\Omega_j) \\ &= \sum_{j=1}^N p_j \times \frac{1 - \left( 1 - \frac{H_j}{H} + \sum_{b \in C(j, \text{root})} \frac{L_b}{H} p_b \right)^{\alpha-1}}{\alpha - 1} \end{aligned}$ | $\begin{aligned} {}^{\alpha}I &= \sum_{j=1}^N p_j ({}^{\alpha}\Omega_j^*) \\ &= \sum_{j=1}^N p_j \left[ \sum_{b \in C(j, \text{root})} L_b \frac{1 - p_b^{\alpha-1}}{\alpha - 1} \right] \end{aligned}$ |
| Species equivalents<sup>c</sup> | ${}^{\alpha}E = \left[ \sum_{j=1}^N p_j \left( 1 - \frac{H_j}{H} + \sum_{b \in C(j, \text{root})} \frac{L_b}{H} p_b \right)^{\alpha-1} \right]^{\frac{1}{1-\alpha}}$ | ${}^{\alpha}E^* = \left[ \sum_{j=1}^N p_j \left( 1 - \frac{H_j}{H} + \sum_{b \in C(j, \text{root})} \frac{L_b}{H} p_b^{\alpha-1} \right) \right]^{\frac{1}{1-\alpha}}$ |

<sup>a</sup> Phylogenetic diversity is here expressed in terms of the length ( $L_b$ ) and relative abundance ( $p_b$ ) associated with each branch $b$ of the phylogenetic tree with species as tips. Notations are identical as in the main text; $H \geq \max_j(H_j)$ . Abundance-based rarity ( ${}^{\alpha}\rho_j$ ) and species diversity ( ${}^{\alpha}S$ ) are here identical as in Table 1.
<sup>b</sup> ${}^{\alpha}Y = {}^{\alpha}K$ and ${}^{\alpha}I = {}^{\alpha}K^*$ (described in Table 1) if ${}^{\alpha}K$ and ${}^{\alpha}K^*$ are applied to $d_{ij}$ , the sum of branch lengths on the shortest path from tip $j$ to its most recent common ancestor with species $i$ . ${}^{\alpha}I$ was first developed by Pavoine et al. (2009) and expressed in terms of species originalities by Kondratyeva et al. (2019).
<sup>c</sup> See Chao et al. (2010) for functions related to ${}^{\alpha}E^*$ .

All these diversity indices can be easily transformed into equivalent numbers of species: the number of evenly and maximally dissimilar species needed to obtain the level of FP-diversity observed in an assemblage (Tables 1 and 2). The functions that transform the diversity indices we discussed here in terms of equivalent numbers of species do not change the way species assemblages are ranked from the least to the most diverse (Appendix A).

## 4. The special case of abundance-free distinctiveness indices

Abundance-free distinctiveness indices, particularly phylogenetic distinctiveness indices, are often used in conservation biology (Isaac et al., 2007; Redding et al., 2014). Imposing equal relative abundances for all species in our framework provides a useful family of such distinctiveness indices where abundance data is discarded as outlined below (see a complete introduction of the family in Appendix B).

In the special case of equal abundance for all species (*p_j_*=1/*N* for all *j*), the phylogenetic distinctiveness index *^α^* Δ* associated with phylogenetic diversity index *^α^I* (Table 2) becomes:

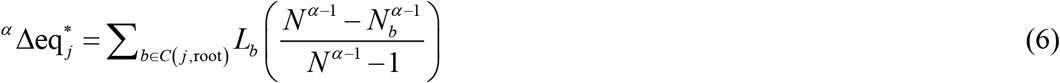

where *N_b_* stands for the number of species that descend from branch *b* and *C*(*j*, root) for the set of branches between species *j*, tip of the phylogenetic tree, and the root of the tree.

*^α^* Δeq* thus provides a parametric alternative to the most widely used index of species distinctiveness named “evolutionary distinctiveness” (*ED*; Isaac et al., 2007) or “Fair-Proportion” (Redding et al., 2014) and whose formula is:

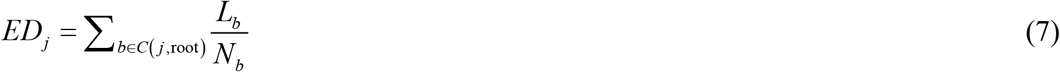

Both *ED* and 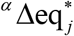 can be seen as the sum, on the shortest path from a tip to root, of the product of a branch length times a decreasing function of the number of species descending from that branch. While by construction, the value of *ED* is dominated by the length of the terminal branch that connects species *j* to the tree (Redding et al., 2014), the parameter *α* in ^*α*^ Δeq* allows controlling the influence of this terminal branch (see Appendix B for more details).

It can be easily shown for example that with α = 0, 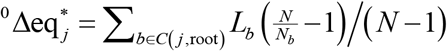 and with α → 1, 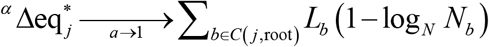 (Appendix B). Another notable case is obtained with α = 2 because

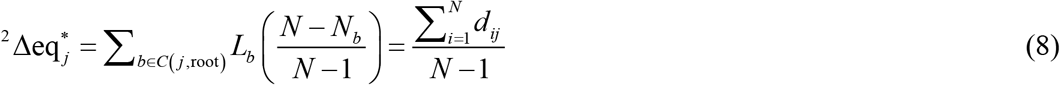

where *d_ij_* is the sum of branch lengths on the shortest path from tip *j* to its most recent common ancestor with species *i*.

Although the diversity indices of Tables 1 and 2 are meaningful only for nonnegative values of *α* (as otherwise rarity function rapidly tends to infinity for low values of *α*), negative values are meaningful for calculating 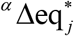. Indeed, 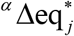 varies between the length, *h_j_*, of the terminal branch that connects *j* to the rest of the tree (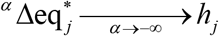, Appendix B), and the distance, 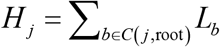, from tip *j* to the root of the tree (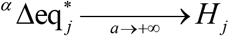, Appendix B). As a consequence, by varying parameter *α* between −∞ and 2 in 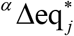 one can obtain a range of strongly connected indices of phylogenetic distinctiveness along a gradient that goes from a strong influence of the terminal branch to the average distance between a species and all others in phylogenetic tree. Index 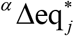 thus responds to the critical need in conservation science to justify and organize indices of evolutionary distinctiveness which currently represent different trade-offs between the unique evolutionary history represented by a species versus its average distance to all other species, trade-offs that may impact on-the-ground decision on conservation priority (Redding et al., 2014). Similarly, using even abundance in the distinctiveness index 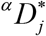 (Table 1) provides a measure of FP-distinctiveness *α Deq*\* whose values vary (by varying ^*α*^) from the smallest FP-dissimilarity between *j* and any other species (*α* → −∞), through the average FP-dissimilarity to all other species (*α* = 2), to the largest FP-dissimilarity between *j* and any other species (*α* → +∞) (Appendix B).

## 5. Worked example

### 5.1. Data

As a case study, we considered changes in bat phylogenetic diversity across a disturbance gradient in the Selva Lacandona of Chiapas, Mexico. Medellín et al. (2000) collected the abundance of bat species in four habitats (rainforests, cacao plantations, old fields and cornfields). We obtained the phylogeny of 34 observed species using a consensus ultrametric tree (function consensus.edge, R package phytools; Revell, 2012) on 9,999 credible birth-death, tip-dated completed trees downloaded from http://vertlife.org/phylosubsets on 2021/02/12 (Upham et al., 2019). For species names, we followed Upham et al. (2019). Branch lengths on the consensus tree were obtained using mean edge length, ignoring credible trees in which the branch is absent (Revell, 2012).

### 5.2. Analyses

We calculated, in each habitat, the phylogenetic diversity using *^α^Y* and *^α^I*, for *α* varying between 0 and 3. We explored then in detail the patterns of phylogenetic diversity in terms of effective originality. Additionally, we analyzed species distinctiveness in the species pool (using indices *ED* and *^α^* Δeq* that both discard abundance data).

### 5.3. Results

In the species pool, for values of *α* of 1, 2 and 3, according to *^α^* Δeq* the two Vespertilionidae species (*Bauerus dubiaquercus* and *Myotis keaysi*) are the most phylogenetically distinct, being the sole members of their clade in the dataset (Fig. 2). The least distinctive species are the two *Artibeus* species as they are in the most species-rich part of the phylogenetic tree, where both terminal and internal branches are numerous and relatively short. For low values of *α*, the distinctiveness is strongly influenced by the length of the terminal branches. This has two main consequences: the species perceived as least distinct are the two *Carollia* species that diverged recently; and the two Vespertilionidae species are no longer perceived as the most distinct, as their terminal branches are shorter than those of *Thyroptera tricolor*, *Pteronotus parnellii* and *Mormoops megalophylla*. The case, *α* = 0 is the closest to *ED* (Pearson correlation = 1). For ^0^ Δeq* and *ED*, *T. tricolor* is the most distinctive followed by the two Vespertilionidae and then *P. parnellii* and *M. megalophylla*; the least distinctive are the *Artibeus* species followed by the *Dermanura* species and then the *Carollia* species. A more complete visualization of variation in *^α^* Δeq* as a function of *α* can be found in video S1.

**Fig. 2.**
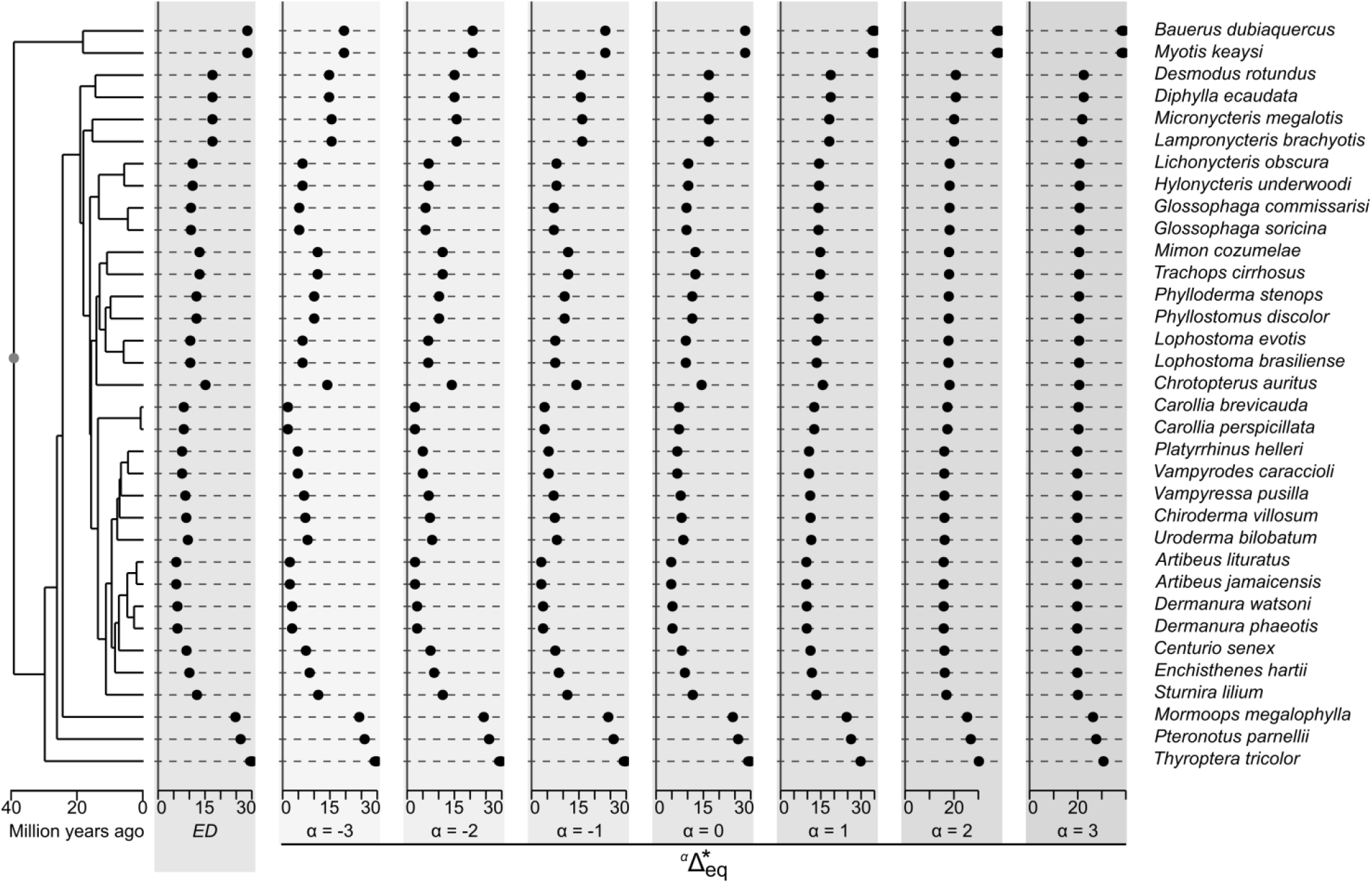
Phylogenetic species distinctiveness. (measured considering the 34 species of the dataset by *ED* and then ^*α*^Δeq*, from *α* = −3 to *α* = 3 with an increment of 1; the grey dot on the phylogenetic tree indicates the root).

Phylogenetic diversity decreases along the disturbance gradient (from rainforest, through cacao plantation and old fields, to cornfields) whatever the value of *α* we considered (from 0 to 3 in Fig. 3) for *^α^I* and for high values of *α* (*α* >= 1.5 in Fig. 3) for *^α^Y*. With *^α^Y*, for low values of *α* (*α* < 1.5 in Fig. 3) the phylogenetic diversity of old fields exceeds that of cacao plantations; and as *α* approaches zero (*α* < 0.5 in Fig. 3) the phylogenetic diversity of old fields even exceeds that of the rainforest. This is because old fields contain both *B. dubiaquercus* and *M. keaysi*, the two Vespertilionidae species with the highest average phylogenetic distance to all other species, while only one of them was observed in the rainforest and in the cacao plantation, and none in the cornfields (Fig. 4). Indeed index *^α^Y*, as *^α^K*, uses the effective originality associated with the quadratic diversity and a parameter *α* that controls the relative importance given to ordinary species compared to effectively original species. For low values of *α*, the influence of the most effectively original species in the measurement of phylogenetic diversity increases. In contrast as shown above, *α* in *^α^I*, as in *^α^K*\*, controls the importance given to abundant compared to rare species, influencing the way the effective originality of a species is perceived (with strong influence of the terminal branch for low values of *α*). Compared to other habitats, cornfields lacked effectively original species (Fig. 4; see also Video S2 for a more complete visualization of variation in species originality *^α^* Ω* as a function of *α*). Phylogenetic diversity in cornfields was thus always the smallest (whatever *α*; and even strongly lower than that of other habitats when *α* approaches zero; Fig. 3). Species with the least effective originality were often either the *Carollia* species or the *Artibeus* species depending on the habitat and the value of *α* considered (Fig. 4). However the relatively high abundance of *P. parnelii* observed in the rainforest, made this species perceived as one of the least effectively original in this habitat when rare species were given high weights in the measurement of phylogenetic diversity (*α* = 0, index *^α^I*, Fig. 4), despite its high distinctiveness (Fig. 2 and 4; see also Video S3 for a more complete visualization of variation in species distinctiveness *^α^* Δ* as a function of *α*).

**Fig. 3.**
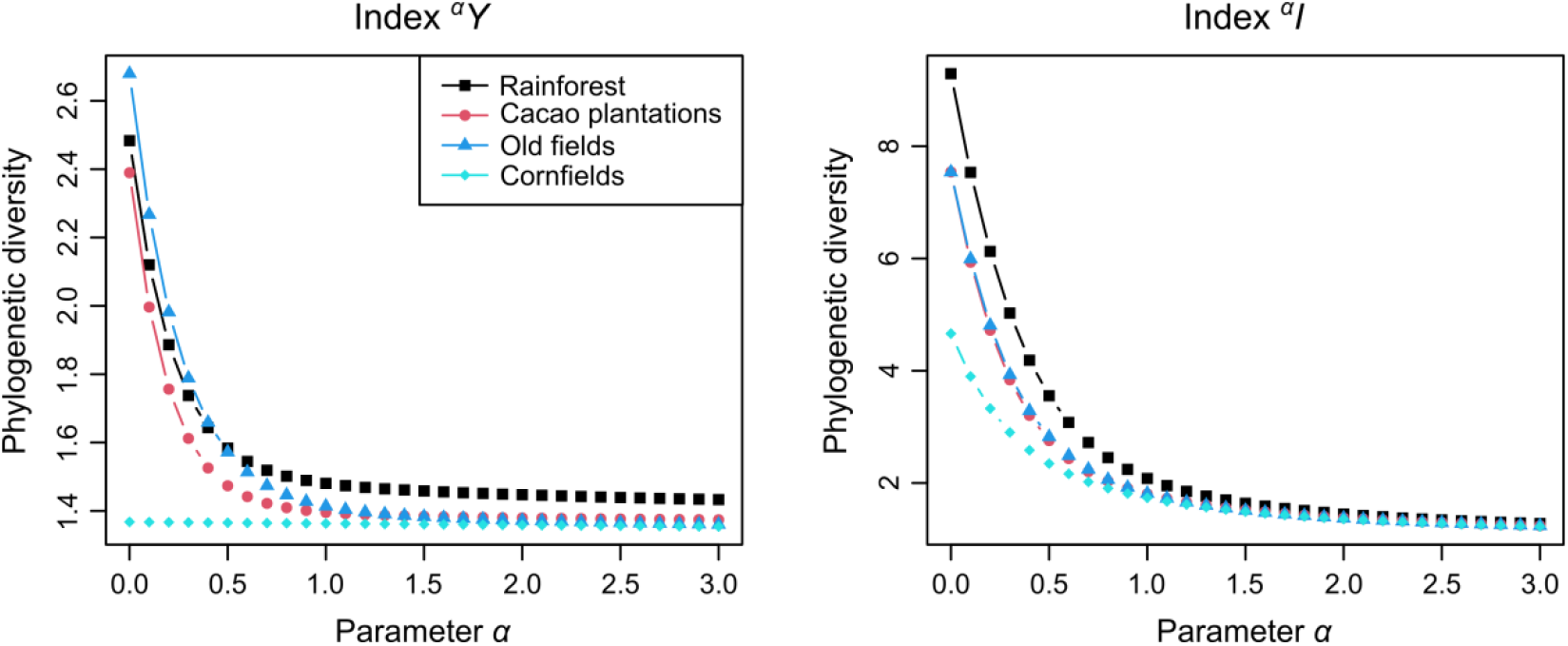
Phylogenetic diversity of the four habitats. (Indices are here expressed as equivalent numbers of species).

**Fig. 4.**
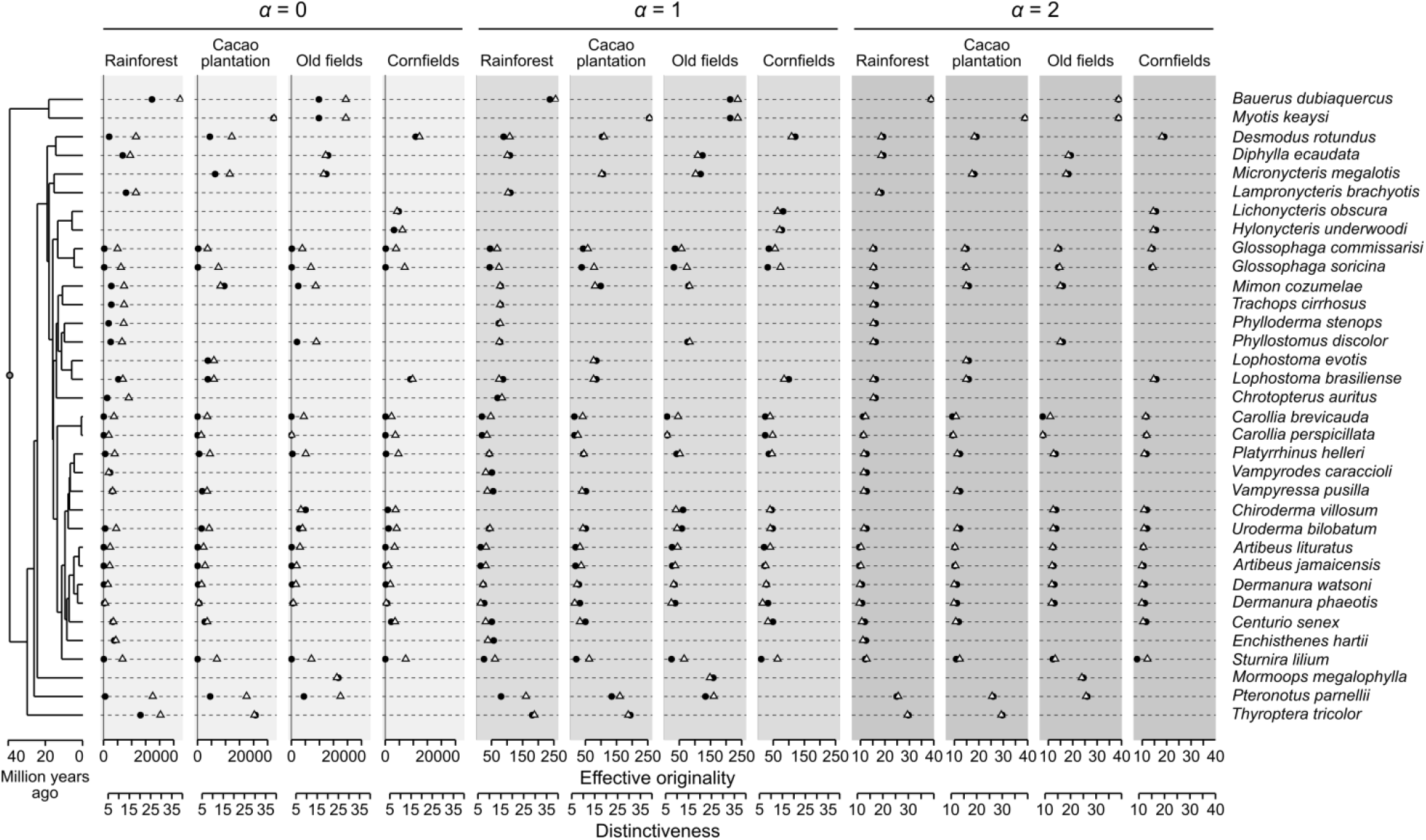
Species effective originality and distinctiveness in rainforest, cacao plantation, old fields, and cornfields. (Effective originality (black circles) and distinctiveness (white triangles) were measured here with indices *^α^*Ω* and *^α^* Δ*, respectively, considering from left to right, *α* = 0, *α* = 1, and *α* = 2. The parameter value *α* = 2 corresponds to the case where measurements for *^α^I*, *^α^Y* and *Q* merge. For the effective originality and distinctiveness measures associated with *^α^Y*, varying the *α* parameter does not change the ranking of species from the least to the most effectively original species, which corresponds here to the ranking obtained with *α* = 2.)

## 6. Discussion

The contribution of a given species to the biodiversity of an assemblage thus depends on its rarity and on the rarity of its FP-characteristics. Starting from Ricotta et al. (2016) measure of species-level originality *K* we have shown that quadratic diversity can be expressed as a mean of effective FP-originality values over all species in an assemblage, and that Faith’s (1992) phylogenetic diversity index and Cadotte et al. (2010) and Kondratyeva et al. (2019) measures of effective phylogenetic originality can be both related to parametric extensions of quadratic diversity. This led us to develop a unified framework (summarized in Tables 1 and 2) where diversity, rarity, distinctiveness and originality measures are intrinsically linked.

The parametric indices developed in this framework allow regulating the importance given to abundant and ordinary species in FP-diversity measures. They include generalizations of the Shannon index (for α → 1) and the Simpson index (for α = 2) to functional and phylogenetic data. By increasing the value of their parameter *α*, the weight given to abundant and ordinary species in the measurement of FP-diversity increases in comparison with the weight given to rare and effectively original species. Low values of *α* may thus indicate regions with high diversity but a diversity that may be threatened by the rarity of the most effectively original species. Increasing *α* may reveal how much phylogenetic diversity depends on these rare species. Low values of the parameter *α* could thus be particularly relevant to obtain biodiversity indicators directed to the preservation of rare and distinct species while maintaining a high level of global diversity (e.g., Hidasi-Neto, Loyola and Cianciaruso, 2015).

In our case study, when effectively original species were given high weights in the measurement of diversity (according to *^α^Y* which evaluates effective originality as a function of the average phylogenetic distance to all individuals observed in the habitat), the old fields had the highest measure of phylogenetic diversity. This was due to the presence of the two species from the Vespertilionidae family. Although distinct in our study area, the Vespertilionidae species represent a large family of bats at a global scale. This illustrates how the measurement of a species’ distinctiveness is dependent on the reference species assemblage, on the data used to characterize species (here phylogeny), and thus on the taxonomic, phylogenetic and spatial scales of a study. A species’ rarity, when measured relatively to the rarity of all other species rather than as an absolute value, is also dependent on these scales. According to the International Union for the Conservation of Nature (IUCN, 2021), all species of our case study are least-concern (i.e., neither threatened nor near threatened) at a global scale with either stable or unknown population trends except *M. megalophylla* that is least-concern but with decreasing population trends, and *B. dubiaquercus* that is near threatened (+ *Vampyressa pusilla* classified as data deficient). Our results showed that *M. megalophylla* and *B. dubiaquercus* are among the most phylogenetically distinct species in our study area. Only one individuals of *M. megalophylla* and three individuals of *B. dubiaquercus* were observed in our case study (out of a total of 2405 bats). *M. megalophylla* was observed in old fields and *B. dubiaquercus* in rainforest and old fields. These scarce occurrences prevented us to evaluate the possible direct effects of agriculture on their population size. However, both species are aerial insectivous species (Rodríguez-Aguilar et al., 2017) and insect-eating bats are known to be affected when high pesticide inputs are used in plantations (e.g., Estrada et al., 2006). Other studies showed that, in addition to cave collapse, cave vandalism, threats on *M. megalophylla* and *B. dubiaquercus* also concern their sensitivity to disturbance and habitat loss (IUCN, 2021). In an urban context in the highlands of Chiapas, it was shown that abundance of *B. dubiaquercus* tends to diminish outside forest; and the activity of both *M. megalophylla* and *B. dubiaquercus* increases with tree density (Rodríguez-Aguilar et al., 2017).

The whole framework can thus be used in ecological studies to reveal the relative contributions of each species to biodiversity, and to depict these contributions in terms of abundance-based rarity, species-level FP-distinctiveness and effective FP-originality. It can be applied from local to global scales provided abundance data are available and the species assemblage is clearly delimited. The framework can be used to evaluate how species contributions to biodiversity vary in space and which evolutionary and/or ecological processes have shaped the biodiversity of an assemblage. Studying a process of invasion for example, the framework could be used across various scales and biomes, to confront Darwin’s naturalization hypothesis, specifying that the taxonomic distance (or phylogenetic and functional distinctiveness) of an invader compared with native species increases the chance of successful installation by limiting competitions, and Darwin’s alternative niche-adaptation hypothesis, for which ecological (or functional) redundancy with natives is expected so that the invader is pre-adapted or tolerant to the local environmental conditions (Darwin, 1859; see, e.g., Park et al., 2020). In this context, weighting originality measures by species abundance is critical to evaluate the potential degree of competition between species individuals and how competition could influence the functional and phylogenetic compositions of an assemblage.

Our framework can be used to evaluate how changes in species contributions may impact both biodiversity levels and ecosystem functioning. Indeed, complementing abundance data with originality values is important in this context. As indicated in the introduction, rare species (of low abundance) may contribute more to ecosystem functioning and ecosystem services than their low abundance would suggest because of their unique functional characteristics (Dee et al., 2019). Depicting how rarity and functional distinctiveness influence measures of biodiversity could thus improve research on the connections between biodiversity and ecosystem functioning.

In the context of the sixth species mass extinction (Ceballos et al., 2015), our framework could be used to evaluate how changes in species contributions may impact ecosystem services, via changes in biodiversity levels. It would allow following changes in species contribution to biodiversity under novel environmental conditions, from local to global planetary-scale change. Relative abundances may change under novel environmental conditions with rare species for example benefiting from the reduction of population size of other species. If these species are functionally rare, an increase in their abundance can considerably increase the functional diversity of an assemblage and modify ecosystem functioning and the connected ecosystem services. The bioprospecting or option value associated to a species notably advocates as a precaution principle to protect species with the aim to give option to discover new uses of these species in the future, especially in medicine. Here functionally and phylogenetically unique species may be considered as disproportionally contributing to bioprospecting value (Dee et al., 2019). Where and when they have low abundance, they might be in need of urgent conservation actions (e.g., the Western Swamp Tortoise cited above, Arnall, 2018; or the Van Gelder’s bat, *B. dubiaquercus*, of our case study). Indeed, if environmental changes inversely lead to the extinction of currently effectively original species with key role in the ecosystem, these changes could yield the biological system to collapse, with potential drastic loss of ecosystem services. Effectively original species may for example be directly threatened when they are increasingly targeted by economic activities because of their combined aspects of rarity (e.g., private collections of rare, distinct specimens; safaris spotting rare, distinct species; e.g., Holden and McDonald-Madden, 2017).

Links between the functional distinctiveness of a species and its abundance-based rarity have for example been observed in European estuarine fish communities, with identified potential consequences on the stability of these communities (Teichert et al., 2017). Links between the functional distinctiveness of a species and its risk of extinction have also sometimes been observed: e.g., among anurans in Ecuador (Menéndes-Guerrero et al., 2020); globally in mammals and birds (Cooke et al., 2020). Species which are rare both in terms of low abundance and phylogenetic distinctiveness have sometimes been found to be threatened. For example, Uchida et al. (2019) observed that, in semi-natural grasslands of south west Japan, low-abundance and phylogenetically distinct species were threatened by land-use intensification, resulting in plant phylogenetic diversity loss. In another context, at a global scale, Sol et al. (2017) found a decrease in abundance or even loss of phylogenetically distinct bird species in highly-urbanized areas compared to the surrounding natural environments.

Our framework thus allows depicting biodiversity in terms of rarity, distinctiveness, and effective originality to better identify how species together contribute to biodiversity levels. It has the potential to improve studies on the mechanisms by which global changes affect biodiversity levels, by identifying which aspect of biodiversity they impact, be it rarity, functional distinctiveness or the global effective originality of some species. Our framework unifies various measures of diversity, distinctiveness and originality, previously scattered in the literature and developed in different context. By developing it, we revealed a parametric family of phylogenetic distinctiveness indices that could complement the most currently used “evolutionary distinctiveness” index (e.g. Isaac et al., 2007; Ibáñez-álamo et al., 2017; Potter 2018; Cooke et al., 2020) whose values are strongly dominated by the independent evolutionary history of a species (length of terminal branch in a phylogenetic tree with the species as tip; Redding et al., 2014). The parametric family indeed allows controlling the degree of influence of this independent evolutionary history on the distinctiveness index. We provide also an equivalent framework for functional distinctiveness. Overall, our framework helps to provide justification, explanation and order when applying a quantitative reasoning to biodiversity, improving then the research on the mechanisms than drive biodiversity changes, and contributing to the development of efficient biodiversity indicators for conservation strategies.

## Data statement

All data and R scripts are currently placed in Appendixes C to F. They will soon be also placed in the adiv package of R.

