## Appendix A for "On the relationships between rarity, uniqueness, distinctiveness, originality and functional/phylogenetic diversity"

**Appendix A. Detailed presentation of the parametric framework, discussion and proofs on some properties of the indices**

**A.1. On the distinctiveness associated with Rao's quadratic entropy**

A community-level measure of expected FP-distinctiveness may be obtained as the weighted average of species-level distinctiveness for the entire assemblage $D = \sum_{j=1}^{N} p_j \times D_j$ . $D$ is high if the most abundant species have the highest FP-distinctiveness. Considering the most extreme scenario, $D$ attains its maximum value when a single species, say $j$, is maximally dissimilar from all others (e.g., for $d_{ij}$ bounded in [0, 1], $d_{ij}$ = 1 for all $i \neq j$ and $d_{ik}$ tending to 0 for all $i,k \neq j$) and dominates in abundance (i.e., $p_j$ tends to 1 and $p_i$ tends to zero for all $i \neq j$). As such, in general, $D$ is not directly interpretable as an index of FP-diversity as it does not increase with the range of different functional traits or phylogenetic positions that all individuals from all species in the assemblage have. However, $D$ is interpretable as an index of FP-diversity in the special case where species always have equal abundance (i.e., $p_j = 1/N \ \forall j$ ). Indeed in that special case $D$ is simply the average dissimilarity between two species (excluding the comparison of a species with itself), usually named in the literature as the mean pairwise distance ($MPD$) in a phylogenetic context: $MPD = \sum_{i=1}^{N} \sum_{j=1}^{N} \frac{d_{ij}}{N(N-1)}$ (e.g. Webb, 2000).

**A.2. On the parametric index $^{\alpha}K$**

Functions $\rho_j$ and $O_j$ associated with Rao's quadratic entropy can be traced back to the same moment of the generalized diversity $^{\alpha}K$ of Ricotta and Szeidl (2006). For $\alpha = 2$, if the species are treated as maximally distinct from each other (i.e., $d_{ij}$ = 1 for all $i \neq j$), the generalized effective FP-originality function $^{\alpha}O_j = (1-\omega_j^{\alpha-1})/(\alpha-1)$ (with $\omega_j = \sum_{j=1}^{N} p_j s_{ij}$ ) reduces to the Simpson rarity $\rho_j = (1-p_j)$, whereas if the species are not treated as maximally distinct from each other, we get the effective FP-originality function of the Rao quadratic diversity $O_j = (1-\omega_j)$ . For any value of $\alpha$, considering that species are treated as maximally distinct from each other, $^{\alpha}O_j$ reduces to a parametric version of the Simpson rarity function $^{\alpha}\rho = \left(1-p_j^{\alpha-1}\right)/\left(\alpha-1\right)$.

For $\alpha$ tending to 1 (Ricotta and Szeidl, 2006),

$$^{\alpha}K \to -\sum_{j=1}^{N} p_j \ln \omega_j$$

This limit differs from that of index $^{\alpha}K^*$ for which

$$^{\alpha}K^* \xrightarrow[\alpha \to 1]{} -\sum_{j=1}^{N} p_j \times \sum_{c=1}^{N-1} u_{c|j} \ln\left(p_{c|j}\right)$$

$^{\alpha}K$ can be also seen as a particular application of a weighted version of the generalized diversity of Patil and Taillie (1982) or equivalently of a parametric generalization of the 'weighted Gini-Simpson index of diversity' developed by Guiasu and Guiasu (2010). Indeed, apart from the Rao quadratic diversity, another diversity index has been proposed as a generalization of the Simpson index by Guiasu and Guiasu (2010, 2012) under the name of 'weighted Gini-Simpson index of diversity':

$$G_w = \sum_{j=1}^{N} w_j p_j (1-p_j) \qquad \text{(A.1)}$$

where the species-specific weights $w_j$ $(j=1,2,...,N)$ are non-negative real numbers which may reflect the ecological importance, rarity, or economic value of the species in a given assemblage (Guiasu and Guiasu 2010). Compared with Rao's quadratic diversity, the $G_w$ index usually does not conform to the requirement proposed by Leinster and Cobbold (2012) that diversity should not change if a given species *j* is replaced by two identical species with the same total abundance of *j* (see section A.6. below). However, it may satisfy this requirement for specific definitions of the $w_j$ as shown below.

If $w_j = D_j$ the weighted Gini-Simpson index becomes Rao's diversity: $Q = \sum_{j=1}^{N} D_j p_j (1-p_j) = \sum_{j=1}^{N} p_j (1-\omega_j)$. That is, Rao's quadratic diversity can be also interpreted as a particular formulation of the Simpson index in which the single-species contributions $p_j(1-p_j)$ to community-level diversity are weighted by their species-level FP-distinctiveness $D_j$.

The 'weighted Gini-Simpson index of diversity' can be further used to relate the generalized diversity of Ricotta and Szeidl (2006) to a weighted version of the generalized diversity of Patil and Taillie (1982) proposed here for the first time:

$${}^{\alpha}G_w = \sum_{j=1}^{N} w_j p_j \times \frac{1-p_j^{\alpha-1}}{\alpha-1} \qquad \text{(A.2)}$$

If the weights $w_j$ in Eq. A.2 are selected as the ratio between two commensurate rarity functions such that $w_j = {}^{\alpha}D_j = (1-\omega_j^{\alpha-1}) \big/ (1-p_j^{\alpha-1})$, Eq. A.2 becomes the generalized diversity of Ricotta and Szeidl (2006) as $\sum_{j=1}^{N} {}^{\alpha}D_j p_j \frac{1-p_j^{\alpha-1}}{\alpha-1} = \sum_{j=1}^{N} p_j \frac{1-\omega_j^{\alpha-1}}{\alpha-1}$, where ${}^{\alpha}D_j$ is the parametric measure of species-level FP-distinctiveness that links the weighted diversity of Patil and Taillie (${}^{\alpha}G_w$) to ${}^{\alpha}K$. For example, for $\alpha$ tending to 1, Eq. A.2 tends to a weighted version of the Shannon entropy (Shannon 1948), first introduced in information theory by Belis and Guiasu (1968): $H_w = -\sum_{j=1}^{N} w_j p_j \ln p_j$. By selecting the values of $w_j$ in $H_w$ as $w_j = (-\ln \omega_j) \big/ (-\ln p_j)$, we obtain $-\sum_{j=1}^{N} p_j \ln \omega_j$, which is the Shannon-like expression of the generalized diversity of Ricotta and Szeidl (2006) for $\alpha$ tending to 1.

### A.3. On the phylogenetic diversity index ${}^{\alpha}Y$

Let $d_{ij}$ be defined as the sum of branch lengths on the shortest path from tip *j* to its most recent common ancestor with species *i*. $d_{ij} = \sum_{b \in C(j, R_{ij})} L_b$, with $R_{ij}$ the most recent common ancestor between species *i* and *j* and $C(j, R_{ij})$ the set of branches between *j* and $R_{ij}$. Let $s_{ij} = 1 - d_{ij} / H$ be the similarity between *i* and *j* with $H \geq \max_{ij}(d_{ij})$. Consider species *j* and

$$^{\alpha}K=\sum_{j=1}^{N}p_j\times\frac{1-\left(\sum_{i=1}^{N}p_i\left(1-\frac{d_{ij}}{H}\right)\right)^{\alpha-1}}{\alpha-1}$$

$$^{\alpha}K=\sum_{j=1}^{N}p_j\times\frac{1-\left(1-\sum_{i=1}^{N}p_i\sum_{b\in C\left(j,\mathrm{R}_{ij}\right)}\frac{L_b}{H}\right)^{\alpha-1}}{\alpha-1}$$

In $\sum_{i=1}^{N}p_i\sum_{b\in C\left(j,\mathrm{R}_{ij}\right)}\frac{L_b}{H}$, each branch on the path from $j$ to root (i.e., in $C(j,\text{ root})$) is multiplied by the sum of relative abundances of all species that do not descend from it, that is to say by 1 minus the sum of relative abundances of the species descending from it:

$$\sum_{i=1}^{N}p_i\sum_{b\in C\left(j,\mathrm{R}_{ij}\right)}\frac{L_b}{H}=\sum_{b\in C\left(j,\mathrm{root}\right)}\frac{L_b}{H}\left(1-p_b\right)=\frac{H_j}{H}-\sum_{b\in C\left(j,\mathrm{root}\right)}\frac{L_b}{H}p_b$$

This yields,

$$^{\alpha}K={}^{\alpha}Y=\sum_{j=1}^{N}p_j\times\frac{1-\left(1-\frac{H_j}{H}+\sum_{b\in C\left(j,\mathrm{root}\right)}\frac{L_b}{H}p_b\right)^{\alpha-1}}{\alpha-1}$$

### A.4. The use of asymmetrical dissimilarities and non-ultrametric trees

$Q$ and $^{\alpha}K$ were originally developed for symmetric dissimilarities between species (i.e., $d_{ij}=d_{ji}$) and $^{\alpha}I$ for ultrametric phylogenetic trees (i.e., trees with constant distance from tip to root). However, our framework is actually still valid if $Q$, $^{\alpha}K$ and $^{\alpha}K^{*}$ are applied to asymmetric dissimilarities (i.e. $d_{ij}$ may be different from $d_{ji}$) and $^{\alpha}Y$ and $^{\alpha}I$ to non-ultrametric phylogenetic trees. The links between the five indices and their associated components of rarity, distinctiveness and originality also still hold with asymmetric dissimilarities and non-ultrametric trees. As far as we are aware, Hendrickson and Ehrlich (1971) developed the first version of what will be named later quadratic diversity by Rao (1982). Hendrickson and Ehrlich (1971) considered as an example of symmetric interspecies differences the extent of niche non-overlap, relative to the combined niche of the species. In this context, asymmetric distances could be envisaged for example if say 80% of the functional niche of a species $i$ overlap with that of species $j$ although the functional niche of species $j$ is larger and only say 10% of its functional niche overlap with that of species $i$. Using non-ultrametric phylogenetic trees may also be useful if the evolutionary rates are not constant among lineages as in that case phylogenetic diversity measures computed from a non-ultrametric tree could yield results different than those expected if they were applied to an ultrametric tree (e.g., Gonzalez, 2010).

### A.5. About equivalent numbers of species.

Consider an observed community of $N$ species characterized by their relative abundances ($p_i$) and FP-dissimilarities between them ($d_{ij}$). Consider also a theoretical community with $E$ species that have equal relative abundances and that have a FP-dissimilarity from each other equal to $d_{\max}$, with $d_{\max}\geq\max_{ij}(d_{ij})$. $d_{\max}$ is considered to be the maximum possible FP-dissimilarities between two species. $d_{\max}$ can be set equal to the maximum observed FP-dissimilarity in the community or in reference to a larger community or to all species of a clade, even those not observed in the community. Transforming a diversity index into species equivalents corresponds to finding $E$ such that the diversity of the theoretical community is equal to that of the observed community.

Applied to $Q=\sum_{i=1}^{N}\sum_{j=1}^{N} p_i p_j d_{ij}$ this gives:

$$\sum_{i=1}^{E}\sum_{j=1, j\neq i}^{E} \frac{1}{E}\frac{1}{E} d_{\max} = \sum_{i=1}^{N}\sum_{j=1}^{N} p_i p_j d_{ij}$$

$$\Leftrightarrow \frac{E-1}{E} d_{\max} = \sum_{i=1}^{N}\sum_{j=1}^{N} p_i p_j d_{ij}$$

$$\Leftrightarrow E = \frac{1}{1-\sum_{i=1}^{N}\sum_{j=1}^{N} p_i p_j \frac{d_{ij}}{d_{\max}}}$$

$$\Leftrightarrow E = \frac{1}{\sum_{i=1}^{N}\sum_{j=1}^{N} p_i p_j \left(1-\frac{d_{ij}}{d_{\max}}\right)}$$

The function that links $E$ with $Q$ is
$E=1/(1-Q/d_{max})$
It is a monotonically increasing function. Thus using $E$ or $Q$ does not change the way assemblages are ranked from the least to the most diverse.

Applied to ${}^{\alpha}K = \sum_{j=1}^{N} p_j \times \frac{1-\left(\sum_{i=1}^{N} p_i s_{ij}\right)^{\alpha-1}}{\alpha-1}$ it gives:

$$\sum_{j=1}^{E} \frac{1}{E} \times \frac{1-\frac{1}{E^{\alpha-1}}}{\alpha-1} = \sum_{j=1}^{N} p_j \times \frac{1-\left(\sum_{i=1}^{N} p_i s_{ij}\right)^{\alpha-1}}{\alpha-1}$$

$$\Leftrightarrow 1-\frac{1}{E^{\alpha-1}} = \sum_{j=1}^{N} p_j \left(1-\left(\sum_{i=1}^{N} p_i s_{ij}\right)^{\alpha-1}\right)$$

$$\Leftrightarrow E = \left[1-\sum_{j=1}^{N} p_j \left(1-\left(\sum_{i=1}^{N} p_i s_{ij}\right)^{\alpha-1}\right)\right]^{\frac{1}{1-\alpha}}$$

$$\Leftrightarrow E = \left[\sum_{j=1}^{N} p_j \left(\sum_{i=1}^{N} p_i s_{ij}\right)^{\alpha-1}\right]^{\frac{1}{1-\alpha}}$$

The function that links this new definition of $E$ with ${}^{\alpha}K$ is
$E=\left(1-(\alpha-1)\left({}^{\alpha}K\right)\right)^{\frac{1}{(1-\alpha)}}$ for $\alpha \geq 0$, $\alpha \neq 1$
and
$E=\exp\left({}^{\alpha}K\right)$ for $\alpha = 1$
Using $E$ or ${}^{\alpha}K$ does not change the way assemblages are ranked from the least to the most diverse.

Applied to ${}^{\alpha}K^{*} = \sum_{j=1}^{N} p_j \times \frac{\sum_{c=1}^{N-1} u_{c|j}\left(1-p_{c|j}^{\alpha-1}\right)}{\alpha-1}$, it yields:

$$\sum_{j=1}^{E} \frac{1}{E} \frac{d_{\max}\left(1-\frac{1}{E^{\alpha-1}}\right)}{\alpha-1} = \sum_{j=1}^{N} p_j \times \frac{\sum_{c=1}^{N-1} u_{c|j}\left(1-p_{c|j}^{\alpha-1}\right)}{\alpha-1}$$

$$\Leftrightarrow d_{\max}\left(1-\frac{1}{E^{\alpha-1}}\right) = \sum_{j=1}^{N} p_j \times \sum_{c=1}^{N-1} u_{c|j}\left(1-p_{c|j}^{\alpha-1}\right)$$

$$\Leftrightarrow \frac{1}{E^{\alpha-1}} = 1-\sum_{j=1}^{N} p_j \times \sum_{c=1}^{N-1} \frac{u_{c|j}}{d_{\max}}\left(1-p_{c|j}^{\alpha-1}\right)$$

$$\Leftrightarrow E = \left[1-\sum_{j=1}^{N} p_j \times \sum_{c=1}^{N-1} \frac{u_{c|j}}{d_{\max}}\left(1-p_{c|j}^{\alpha-1}\right)\right]^{\frac{1}{1-\alpha}}$$

$$\Leftrightarrow E = \left[\sum_{j=1}^{N} p_j \left(1-\sum_{c=1}^{N-1} \frac{u_{c|j}}{d_{\max}} + \sum_{c=1}^{N-1} \frac{u_{c|j}}{d_{\max}} p_{c|j}^{\alpha-1}\right)\right]^{\frac{1}{1-\alpha}}$$

$$\Leftrightarrow E = \left[\sum_{j=1}^{N} p_j \left(1-\frac{\max_i d_{ij}}{d_{\max}} + \sum_{c=1}^{N-1} \frac{u_{c|j}}{d_{\max}} p_{c|j}^{\alpha-1}\right)\right]^{\frac{1}{1-\alpha}}$$

The function that links this new definition of $E$ with ${}^{\alpha}K^{*}$ is

$$E = \left(1-\left(\alpha-1\right)\frac{{}^{\alpha}K^{*}}{d_{\max}}\right)^{\frac{1}{(1-\alpha)}}$$

Using $E$ or ${}^{\alpha}K^{*}$ does not change the way assemblages are ranked from the least to the most diverse.

Similarly with a phylogenetic tree, the theoretical community is considered to be composed of $E$ evenly abundant species that are located at the tips of a star-shaped phylogenetic tree, with a single internal node (the root) and branches that connect tips directly to this node with a length of $H$ (with $H \geq H_j$, the sum of branch length on the path from tip $j$ to root in the real tree). As in the main text, $C(j$, root) is the set of nodes between species $j$ and the root of the phylogenetic tree; $b$ is a branch of the tree, $L_b$ its length, and $p_b$ the sum of relative abundance of all species descending from it.

With index ${}^{\alpha}Y$, it yields

$$\sum_{j=1}^{E} \frac{1}{E} \times \frac{1-\frac{1}{E^{\alpha-1}}}{\alpha-1} = \sum_{j=1}^{N} p_j \times \frac{1-\left(1-\frac{H_j}{H}+\sum_{b\in C(j,\text{root})} \frac{L_b}{H} p_b\right)^{\alpha-1}}{\alpha-1}$$

$$\Leftrightarrow 1-\frac{1}{E^{\alpha-1}} = \sum_{j=1}^{N} p_j \left(1-\left(1-\frac{H_j}{H}+\sum_{b\in C(j,\text{root})} \frac{L_b}{H} p_b\right)^{\alpha-1}\right)$$

$$\Leftrightarrow \frac{1}{E^{\alpha-1}} = 1-\sum_{j=1}^{N} p_j \left(1-\left(1-\frac{H_j}{H}+\sum_{b\in C(j,\text{root})} \frac{L_b}{H} p_b\right)^{\alpha-1}\right)$$

$$\Leftrightarrow E=\left[1-\sum_{j=1}^{N} p_j\left(1-\left(1-\tfrac{H_j}{H}+\sum_{b\in C(j,\text{root})}\tfrac{L_b}{H}\,p_b\right)^{\alpha-1}\right)\right]^{\frac{1}{1-\alpha}}$$

$$\Leftrightarrow E=\left[\sum_{j=1}^{N} p_j\left(1-\tfrac{H_j}{H}+\sum_{b\in C(j,\text{root})}\tfrac{L_b}{H}\,p_b\right)^{\alpha-1}\right]^{\frac{1}{1-\alpha}}$$

With ${}^{\alpha}I=\sum_{j=1}^{N} p_j\left[\sum_{b\in C(j,\text{root})} L_b\,\dfrac{1-p_b^{\alpha-1}}{\alpha-1}\right]$, it yields

$$\sum_{j=1}^{E}\frac{1}{E}H\,\frac{1-\frac{1}{E^{\alpha-1}}}{\alpha-1}=\sum_{j=1}^{N} p_j\left[\sum_{b\in C(j,\text{root})} L_b\,\frac{1-p_b^{\alpha-1}}{\alpha-1}\right]$$

$$\Leftrightarrow H\left(1-\tfrac{1}{E^{\alpha-1}}\right)=\sum_{j=1}^{N} p_j\left[\sum_{b\in C(j,\text{root})} L_b\left(1-p_b^{\alpha-1}\right)\right]$$

$$\Leftrightarrow 1-\tfrac{1}{E^{\alpha-1}}=\sum_{j=1}^{N} p_j\left[\sum_{b\in C(j,\text{root})}\frac{L_b}{H}\left(1-p_b^{\alpha-1}\right)\right]$$

$$\Leftrightarrow \tfrac{1}{E^{\alpha-1}}=1-\sum_{j=1}^{N} p_j\left[\sum_{b\in C(j,\text{root})}\frac{L_b}{H}\left(1-p_b^{\alpha-1}\right)\right]$$

$$\Leftrightarrow E=\left[1-\sum_{j=1}^{N} p_j\left(\sum_{b\in C(j,\text{root})}\frac{L_b}{H}\left(1-p_b^{\alpha-1}\right)\right)\right]^{\frac{1}{1-\alpha}}$$

$$\Leftrightarrow E=\left[1-\sum_{j=1}^{N} p_j\frac{H_j}{H}+\sum_{j=1}^{N} p_j\sum_{b\in C(j,\text{root})}\frac{L_b}{H}\,p_b^{\alpha-1}\right]^{\frac{1}{1-\alpha}}$$

$$\Leftrightarrow E=\left[\sum_{j=1}^{N} p_j\left(1-\frac{H_j}{H}+\sum_{b\in C(j,\text{root})}\frac{L_b}{H}\,p_b^{\alpha-1}\right)\right]^{\frac{1}{1-\alpha}}$$

As for ${}^{\alpha}K^{*}$, ${}^{\alpha}K^{*}$, the functions that transform ${}^{\alpha}Y$ and ${}^{\alpha}I$ in equivalent numbers of species are monotonically increasing meaning that applying them does not change the way assemblages are ranked from the least to the most diverse.

**A.6. On species splitting**

Leinster and Cobbold (2012) advocated the development of FP-diversity indices that respect the following property: diversity should not change if a given species *j* is replaced by two identical species *k* and *l* with the same total abundance of *j* (i.e., $p_j = p_k + p_l$). Rao quadratic diversity fulfills this property (Leinster and Cobbold, 2012).

Like for quadratic diversity, the parametric generalizations ${}^{\alpha}K$, ${}^{\alpha}K^{*}$, ${}^{\alpha}Y$ and ${}^{\alpha}I$ also fulfill the requirement that diversity should not change if a given species *j* is replaced by two identical species with the same total abundance of *j*.

Proof that ${}^{\alpha}K$ is unchanged if a species is replaced by two identical species with the same total abundance:

$$^{\alpha}K = \sum_{j=1}^{N} p_j \times \frac{1-\left(\sum_{i=1}^{N} p_i s_{ij}\right)^{\alpha-1}}{\alpha-1}$$

Consider that any species $j$ is split in two identical species $k$ and $l$ with $s_{kj} = s_{lj} = s_{kl} = 1$, $s_{ij} = s_{ik} = s_{il}$ for all $i$ and $p_j = p_k + p_l$. Let $C_N$ be the set of species that includes species $j$ but not $k$ and $l$ and $C_{N+1}$ the alternative set of $N$+1 species that includes species $k$ and $l$ but not $j$.

Because

$$\sum_{i\in C_N} p_i s_{ij} = \sum_{i\in C_N, i\neq j} p_i s_{ij} + p_j s_{jj}$$
$$= \sum_{i\in C_{N+1}, i\neq k,l} p_i s_{ik} + p_k s_{kk} + p_l s_{lk} = \sum_{i\in C_{N+1}} p_i s_{ik}$$
$$= \sum_{i\in C_{N+1}, i\neq k,l} p_i s_{il} + p_k s_{kl} + p_l s_{ll} = \sum_{i\in C_{N+1}} p_i s_{il}$$

we have

$$\sum_{i\in C_N} p_i s_{ij} = \sum_{i\in C_{N+1}} p_i s_{ik} = \sum_{i\in C_{N+1}} p_i s_{il}$$

and

$$p_j \times \frac{1-\left(\sum_{i\in C_N} p_i s_{ij}\right)^{\alpha-1}}{\alpha-1} = p_k \times \frac{1-\left(\sum_{i\in C_{N+1}} p_i s_{ik}\right)^{\alpha-1}}{\alpha-1} + p_l \times \frac{1-\left(\sum_{i\in C_{N+1}} p_i s_{il}\right)^{\alpha-1}}{\alpha-1}$$

In addition for any species $h \neq j$,

$$\sum_{i\in C_N} p_i s_{ih} = \sum_{i\in C_N, i\neq j} p_i s_{ih} + p_j s_{jh}$$
$$= \sum_{i\in C_{N+1}, i\neq k,l} p_i s_{ih} + p_k s_{kh} + p_l s_{lh}$$
$$= \sum_{i\in C_N +1} p_i s_{ih}$$

and thus

$$p_h \times \frac{1-\left(\sum_{i\in C_N} p_i s_{ih}\right)^{\alpha-1}}{\alpha-1} = p_h \times \frac{1-\left(\sum_{i\in C_N +1} p_i s_{ih}\right)^{\alpha-1}}{\alpha-1}$$

□

Proof that $^{\alpha}K^{*}$ is unchanged if a species is replaced by two identical species with the same total abundance:

$$^{\alpha}K^{*} = \sum_{j=1}^{N} p_j \times \frac{\sum_{c=1}^{N-1} u_{c|j}\left(1-p_{c|j}^{\alpha-1}\right)}{\alpha-1}$$

Consider that species $j$ is split in two identical species $k$ and $l$ with $d_{kj} = d_{lj} = d_{kl} = 0$, $d_{ij} = d_{ik} = d_{il}$ for all $i$ and $p_j = p_k + p_l$. As above, let $C_N$ be the set of species that includes species $j$ but not $k$ and $l$ and $C_{N+1}$ the alternative set of $N$+1 species that includes species $k$ and $l$ but not $j$.

Then, for any $c$, $u_{c|j}$ in $C_N$ is equal to $u_{c+1|k} = u_{c+1|l}$ in $C_{N+1}$ because $l$ can be defined as the most similar species to $k$ ($1|k = l$) and similarly $1|l = k$ . As $k$ and $l$ are similar, $u_{1|k} = u_{1|l} = 0$. For any $c > 1$, $p_{c-1|j} = p_{c|k} = p_{c|l}$ and $p_{1|j} = p_j = p_k + p_l = p_{1|k} + p_{2|k} = p_{1|l} + p_{2|l}$. This yields

$$\sum_{c=1}^{N-1} u_{c|j}\left(1-p_{c|j}^{\alpha-1}\right)=\sum_{c=1}^{N-1} u_{c+1|k}\left(1-p_{c+1|k}^{\alpha-1}\right)$$
$$=\sum_{c=2}^{N} u_{c|k}\left(1-p_{c|k}^{\alpha-1}\right)$$
$$=\sum_{c=2}^{N} u_{c|k}\left(1-p_{c|k}^{\alpha-1}\right)+u_{1|k}\left(1-p_{1|k}^{\alpha-1}\right)$$
$$=\sum_{c=1}^{N} u_{c|k}\left(1-p_{c|k}^{\alpha-1}\right)$$

similarly,

$$\sum_{c=1}^{N-1} u_{c|j}\left(1-p_{c|j}^{\alpha-1}\right)=\sum_{c=1}^{N} u_{c|l}\left(1-p_{c|l}^{\alpha-1}\right)$$

which yields

$$p_j \times \frac{\sum_{c=1}^{N-1} u_{c|j}\left(1-p_{c|j}^{\alpha-1}\right)}{\alpha-1}=p_k \times \frac{\sum_{c=1}^{N-1} u_{c|k}\left(1-p_{c|k}^{\alpha-1}\right)}{\alpha-1}+p_l \times \frac{\sum_{c=1}^{N-1} u_{c|l}\left(1-p_{c|l}^{\alpha-1}\right)}{\alpha-1}$$

In addition for any species $h \neq j$, say that in $C_N$, $j$ is $n^{\text{th}}$ most similar species to $h$. Then in $C_{N+1}$, we can set $k$ and $l$ the $n^{\text{th}}$ and $n+1^{\text{th}}$ most similar species to $h$, respectively, as $d_{kl}=0$. Also, for $c < n+1$ $u_{c|h}$ and $p_{c|h}$ in $C_N$ are equal to $u_{c|h}$ and $p_{c|h}$ in $C_{N+1}$, respectively, for $c > n+1$ $u_{c|h}$ and $p_{c|h}$ in $C_N$ are equal to $u_{c+1|h}$ and $p_{c+1|h}$ in $C_{N+1}$, respectively, and $u_{n+1|h}=0$ in $C_{N+1}$. This implies that

$$p_h \times \frac{\sum_{c=1}^{N-1} u_{c|h}\left(1-p_{c|h}^{\alpha-1}\right)}{\alpha-1} \text{ in } C_N$$

is equal to

$$p_h \times \frac{\sum_{c=1}^{N} u_{c|h}\left(1-p_{c|h}^{\alpha-1}\right)}{\alpha-1} \text{ in } C_{N+1}$$

□

Proof that ${}^{a}Y$ is unchanged if a species is replaced by two identical species with the same total abundance:

$${}^{\alpha}Y=\sum_{j=1}^{N} p_j \times \frac{1-\left(\sum_{b\in C(j,\text{root})} \frac{L_b}{H} p_b\right)^{\alpha-1}}{\alpha-1}$$

Consider that species $j$ is split in two identical species $k$ and $l$. Then $C(j$, root)=$C(k$, root)=$C(l$, root) meaning that, as they are considered identical, the three species would theoretically be located on the same terminal node (tip) in the phylogenetic tree. Because $p_j=p_k+p_l$, replacing species $j$ by species $k$ and $l$ leave the $p_b$ values unchanged for all $b \in C(j$, root) (equivalently $C(k$, root), equivalently $C(l$, root)) and also for all other branches of the phylogenetic tree.

□

Proof that ${}^{a}I$ is unchanged if a species is replaced by two identical species with the same total abundance:

$${}^{a}I=\sum_{j=1}^{N} p_j\left[\sum_{b\in C(j,\text{root})} L_b \frac{1-p_b^{a-1}}{a-1}\right]$$

Consider the same notations as in the proof above. Then

$$p_j \left[ \sum_{b \in C(j,\mathrm{root})} L_b \frac{1-p_b^{a-1}}{a-1} \right]$$
$$= p_k \left[ \sum_{b \in C(k,\mathrm{root})} L_b \frac{1-p_b^{a-1}}{a-1} \right] + p_l \left[ \sum_{b \in C(l,\mathrm{root})} L_b \frac{1-p_b^{a-1}}{a-1} \right]$$

□
