## Appendix B for "On the relationships between rarity, uniqueness, distinctiveness, originality and functional/phylogenetic diversity"

**Appendix B. Distinctiveness values in the special case of even abundances for all species.**

In the special case of even abundances for all species, Rao's quadratic diversity is

$$Q\text{eq} = \sum_{i=1}^{N}\sum_{j=1}^{N}\frac{1}{N}\frac{1}{N}d_{ij}$$

the associated rarity and originality indices become, respectively,

$$\rho\text{eq} = \frac{N-1}{N}$$

and

$$O\text{eq}_j = \sum_{i=1}^{N}\frac{1}{N}d_{ij}$$

yielding to the following associated distinctiveness index:

$$D\text{eq}_j = \sum_{i\neq j}^{N}\frac{1}{N-1}d_{ij}$$

The FD-distinctiveness values are given in Tables B.1 and B.2 for the parametric extensions of the quadratic diversity.

Table B.1 Functional distinctiveness indices associated with parametric extensions of Rao's quadratic diversity in the special case of even abundances for all species. Distinctiveness is expressed in terms of dissimilarities ($d_{ij} \geq 0$) or similarities ($s_{ij} = 1 - d_{ij}$, in which case $0 \leq d_{ij} \leq 1$) between species; Notations are identical as those in the main text).

| | First parametric extension (functional approach) | Second parametric extension (functional approach) |
|---|---|---|
| | $^{\alpha}\mathrm{Deq}_j = \dfrac{N^{\alpha-1} - \left(\sum_{i=1}^{N} s_{ij}\right)^{\alpha-1}}{N^{\alpha-1} - 1}$ | $^{\alpha}\mathrm{Deq}_j^* = \dfrac{\sum_{c=1}^{N-1} u_{c\|j}\left(N^{\alpha-1} - c^{\alpha-1}\right)}{\left(N^{\alpha-1} - 1\right)}$ |
| $\alpha$→-∞ | $^{\alpha}\mathrm{Deq}_j \xrightarrow[\alpha\to-\infty]{} 0$ | $^{\alpha}\mathrm{Deq}_j^* = \xrightarrow[\alpha\to-\infty]{} \min_{i\neq j} d_{ij}$ |
| $\alpha$=0 | $^{0}\mathrm{Deq}_j = \dfrac{1/\left(\sum_{i=1}^{N} \frac{1}{N} s_{ij}\right) - 1}{N-1}$ | $^{0}\mathrm{Deq}_j^* = \dfrac{\sum_{c=1}^{N-1} u_{c\|j}\left(N/c - 1\right)}{\left(N-1\right)}$ |
| $\alpha$→1 | $^{\alpha}\mathrm{Deq}_j \xrightarrow[\alpha\to 1]{} 1 - \log_N\left(\sum_{i=1}^{N} s_{ij}\right)$ | $^{\alpha}\mathrm{Deq}_j^* \xrightarrow[\alpha\to 1]{} \sum_{c=1}^{N-1} u_{c\|j}\left(1 - \log_N(c)\right)$ |
| $\alpha$=2 | $^{2}\mathrm{Deq}_j = \sum_{i\neq j}^{N} \dfrac{1}{N-1}\left(1 - s_{ij}\right)$ <br> $= \sum_{i\neq j}^{N} \dfrac{1}{N-1} d_{ij}$ | $^{2}\mathrm{Deq}_j^* = \dfrac{\sum_{c=1}^{N-1} u_{c\|j}\left(N - c\right)}{\left(N-1\right)}$ <br> $= \sum_{i\neq j}^{N} \dfrac{1}{N-1} d_{ij}$ |
| $\alpha$→+∞* | $^{\alpha}\mathrm{Deq}_j \xrightarrow[\alpha\to+\infty]{} 1$ | $^{\alpha}\mathrm{Deq}_j^* = \xrightarrow[\alpha\to+\infty]{} \max_{i\neq j} d_{ij}$ |

* case of all species similar ( $s_{ij} = 1\ \forall i, j$ , or all species positioned on the same tip in the phylogenetic tree) excluded.

Table B.2 Phylogenetic distinctiveness indices associated with parametric extensions of Rao's quadratic diversity in the special case of even abundances for all species. Distinctiveness is expressed in terms of the branch lengths ($L_b$) of a phylogenetic tree with species as tips, the number of species descending from branch $b$ ($N_b$), $C(j$, root) the set of branches from j to the root of the tree, $H_j = \sum_{b\in C(j,\text{root})} L_b$ , $h_j$ the length of the terminal branch that connects species $j$ to the rest of the tree and $H$ a value such that $H \geq \max_j H_j$ (Notations are identical as those in the main text).

| | First parametric extension (phylogenetic approach) | Third parametric extension (phylogenetic approach) |
|---|---|---|
| | ${}^{\alpha}\Delta\text{eq}_j = \dfrac{N^{\alpha-1} - \left(\sum_{b\in C(j,\text{root})} \frac{L_b}{H} N_b\right)^{\alpha-1}}{N^{\alpha-1}-1}$ | ${}^{\alpha}\Delta\text{eq}^*_j = \sum_{b\in C(j,\text{root})} L_b \left(\dfrac{N^{\alpha-1} - N_b^{\alpha-1}}{N^{\alpha-1}-1}\right)$ |
| $\alpha$→-∞ | ${}^{\alpha}\Delta\text{eq}_j \xrightarrow[\alpha\to-\infty]{} 0$ | ${}^{\alpha}\Delta\text{eq}^*_j \xrightarrow[\alpha\to-\infty]{} h_j$ |
| $\alpha$=0 | ${}^{0}\Delta\text{eq}_j == \dfrac{1/\left(\sum_{b\in C(j,\text{root})} \frac{L_b}{H}\frac{N_b}{N}\right)-1}{N-1}$ | ${}^{0}\Delta\text{eq}^*_j = \sum_{b\in C(j,\text{root})} L_b \left(\dfrac{\frac{N}{N_b}-1}{N-1}\right)$ |
| $\alpha$→1 | ${}^{\alpha}\Delta\text{eq}_j \xrightarrow[\alpha\to 1]{} 1-\log_N\left(\sum_{b\in C(j,\text{root})} \frac{L_b}{H} N_b\right)$ | ${}^{\alpha}\Delta\text{eq}^*_j \xrightarrow[\alpha\to 1]{} \sum_{b\in C(j,\text{root})} L_b\left(1-\log_N N_b\right)$ |
| $\alpha$=2* | ${}^{2}\Delta\text{eq}_j = \dfrac{N-\sum_{b\in C(j,\text{root})} \frac{L_b}{H} N_b}{N-1}$ <br> $= \dfrac{H-H_j}{H} + \sum_{i\neq j}^{N} \dfrac{1}{N-1}\dfrac{H-H_j+\delta_{ij}}{H}$ | ${}^{2}\Delta\text{eq}^*_j = \sum_{b\in C(j,\text{root})} L_b \left(\dfrac{N-N_b}{N-1}\right)$ <br> $= \sum_{i\neq j}^{N} \dfrac{1}{N-1}\delta_{ij}$ |
| $\alpha$→+∞** | ${}^{\alpha}\Delta\text{eq}_j \xrightarrow[\alpha\to+\infty]{} 1$ | ${}^{\alpha}\Delta\text{eq}^*_j \xrightarrow[\alpha\to+\infty]{} H_j$ |

* $\delta_{ij}$ , as in the main text, is the sum of branch lengths on the shortest path from tip $j$ to its most recent common ancestor with species $i$. For ultrametric trees $H_j$ is constant over j. If $H$ = Hj, then ${}^{2}\Delta\text{eq}_j = \sum_{i\neq j}^{N} \dfrac{1}{N-1}\dfrac{\delta_{ij}}{\max_i \delta_{ij}}$ .

** case of all species similar (i.e. all species positioned on the same tip in the phylogenetic tree) excluded.

Proofs for the limits:

If species are not all similar, because $\sum_{i=1}^{N} s_{ij} < N$ , then when $\alpha \to +\infty$

$$N^{\alpha-1} - \left(\sum_{i=1}^{N} s_{ij}\right)^{\alpha-1} \sim N^{\alpha-1} \text{ and } N^{\alpha-1} - 1 \sim N^{\alpha-1}$$

yielding

$$\frac{N^{\alpha-1} - \left(\sum_{i=1}^{N} s_{ij}\right)^{\alpha-1}}{N^{\alpha-1} - 1} \xrightarrow[\alpha \to +\infty]{} 1$$

Similarly, because $N_b < N$ ,

$$\frac{N^{\alpha-1} - N_b^{\alpha-1}}{N^{\alpha-1} - 1} \xrightarrow[\alpha \to +\infty]{} 1$$

yielding,

$${}^{\alpha}\Delta\mathrm{eq}_j \xrightarrow[\alpha \to +\infty]{} H_j = \sum_{b \in C(j,\mathrm{root})} L_b$$

The same reasoning yields

$$\frac{\left(N^{\alpha-1} - c^{\alpha-1}\right)}{\left(N^{\alpha-1} - 1\right)} \xrightarrow[\alpha \to +\infty]{} 1$$

and thus to

$$ {}^{\alpha}D\mathrm{eq}_j^* \xrightarrow[\alpha\to+\infty]{} \sum_{c=1}^{N-1} u_{c|j} = \sum_{c=1}^{N-1}\left(d_{c|j,j} - d_{c-1|j,j}\right) = d_{N-1|j,j} - d_{0|j,j} = \max_{i\neq j} d_{ij} $$

Note that in all cases if species are all similar then their distinctiveness is null for all values of $\alpha$.

$$ \frac{1-\left(p_j\right)^{\alpha-1}}{\alpha-1} \xrightarrow[\alpha\to1]{} \ln\left(1/p_j\right) $$ (Patil and Taillie 1982)

and

$$ \frac{1-\left(\sum_{i=1}^{N} p_i s_{ij}\right)^{\alpha-1}}{\alpha-1} \xrightarrow[\alpha\to1]{} \ln\left(1/\sum_{i=1}^{N} p_i s_{ij}\right) $$ (Ricotta and Szeidl 2006)

yields

$$ {}^{1}D\mathrm{eq} = \frac{\ln\left(N/\sum_{i=1}^{N} s_{ij}\right)}{\ln\left(N\right)} = 1-\log_N\left(\sum_{i=1}^{N} s_{ij}\right) $$

$$ {}^{\alpha}\Delta\mathrm{eq}_j = \sum_{b\in C(j,\mathrm{root})} L_b\left(\frac{N^{\alpha-1}-N_b^{\alpha-1}}{N^{\alpha-1}-1}\right) $$

For the phylogenetic tree, we reasonably consider that all branches support more than one species except the terminal branches. In this summation for the equation of ${}^{\alpha}\Delta\mathrm{eq}_j$, $N_b = 1$ thus only when $b$ is the terminal branch that connects species $j$ to the rest of the tree. For all other branches, $N_b > 1$. To highlight this point, ${}^{\alpha}\Delta\mathrm{eq}_j$ can be rewritten as

$$ {}^{\alpha}\Delta\mathrm{eq}_j = h_j + \sum\nolimits_{b\in C(A_j,\mathrm{root})} L_b \left( \frac{N^{\alpha-1} - N_b^{\alpha-1}}{N^{\alpha-1} - 1} \right) $$

where $A_j$ is the most recent ancestor of species *j*. When $\alpha$ tends to -∞, both $N^{\alpha-1}$ and $N_b^{\alpha-1}$ tend to 0 while $N^{\alpha-1}-1$ tends to -1. This yields

$$ \sum\nolimits_{b\in C(A_j,\mathrm{root})} L_b \left( \frac{N^{\alpha-1} - N_b^{\alpha-1}}{N^{\alpha-1} - 1} \right) \xrightarrow[\alpha\to-\infty]{} 0 $$

and thus ${}^{\alpha}\Delta\mathrm{eq}_j \xrightarrow[\alpha\to-\infty]{} h_j$.

Knowing that ${}^{\alpha}D\mathrm{eq}_j^* = \frac{\sum_{c=1}^{N-1} u_{c|j}\left(N^{\alpha-1} - c^{\alpha-1}\right)}{\left(N^{\alpha-1}-1\right)} = u_{1|j} + \frac{\sum_{c=2}^{N-1} u_{c|j}\left(N^{\alpha-1} - c^{\alpha-1}\right)}{\left(N^{\alpha-1}-1\right)}$

$$ \frac{\sum_{c=2}^{N-1} u_{c|j}\left(N^{\alpha-1} - c^{\alpha-1}\right)}{\left(N^{\alpha-1}-1\right)} \xrightarrow[\alpha\to-\infty]{} 0 $$

and

$$ u_{1|j} = \min_{i\neq j} d_{ij}, $$

$$ {}^{\alpha}D\mathrm{eq}_j^* = \xrightarrow[\alpha\to-\infty]{} \min_{i\neq j} d_{ij}. $$

The same reasoning can be applied to ${}^{\alpha}D\mathrm{eq}_j = \frac{N^{\alpha-1} - \left(\sum_{i=1}^{N} s_{ij}\right)^{\alpha-1}}{N^{\alpha-1}-1}$ yielding ${}^{\alpha}D\mathrm{eq}_j \xrightarrow[\alpha\to-\infty]{} 0$.

$$ {}^{\alpha}\Delta\mathrm{eq}_j = \sum\nolimits_{b\in C(j,\mathrm{root})} L_b \left( \frac{1-\left(\frac{N_b}{N}\right)^{\alpha-1}}{1-\frac{1}{N^{\alpha-1}}} \right) $$

Because $N_b < N$,

$$ 1-\left(\frac{N_b}{N}\right)^{\alpha-1} \xrightarrow[\alpha\to 1]{} \ln\left(\frac{N_b}{N}\right) $$

$$ 1-\frac{1}{N^{\alpha-1}} \xrightarrow[\alpha\to 1]{} \ln\left(\frac{1}{N}\right) $$

yielding

$$ {}^{\alpha}\Delta\mathrm{eq}_j \xrightarrow[\alpha\to 1]{} \sum\nolimits_{b\in C(j,\mathrm{root})} L_b \left(1-\log_N N_b\right) $$

As $c < N$, the same reasoning yields ${}^{\alpha}D\mathrm{eq}_j^{*} \xrightarrow[\alpha\to 1]{} \sum\nolimits_{c=1}^{N-1} u_{c|j} \left(1-\log_N(c)\right)$.
