## Appendix C for "On the relationships between rarity, uniqueness, distinctiveness, originality and functional/phylogenetic diversity"

#### Appendix C. Manual for R scripts

##### C.1. Parametric Indices of Functional and Phylogenetic Diversity

Function `FPdivparam` calculates functional or phylogenetic (FP-)diversity in communities using parametric indices ${}^{\alpha}K$, ${}^{\alpha}K^*$, ${}^{\alpha}Y$ and ${}^{\alpha}I$ discussed and developed in the main text.
The function `plot.FPdivparam` plots the results of function `FPivparam`.

***Usage:***

```
FPdivparam(comm, disORtree, method = c("KY", "KstarI"), palpha = 2,
   equivalent = FALSE, option = c("asymmetric", "symmetric"),
   dmax = NULL, tol = 1e-8)

plot.FPdivparam(x, legend = TRUE,
   legendposi = "topright", axisLABEL = "FP-diversity",
   type="b", col = if(is.numeric(x)) NULL
   else sample(colors(distinct = TRUE), nrow(x$div)),
   lty = if(is.numeric(x)) NULL else rep(1, nrow(x$div)),
   pch = if(is.numeric(x)) NULL else rep(19, nrow(x$div)),
   ...)
```

***Arguments:***

`comm` a data frame or a matrix typically with communities as rows, species as columns and abundance as entry. Species should be labelled as in object disORtree.

`disORtree` an object inheriting the class `dist`, giving the FP-dissimilarities between species, or inheriting the class `phylo` (see package ape, Paradis and Schliep, 2019), `phylo4` (see package `phylobase, R Hackathon et al., 2020`), or `hclust`, where species are tips.

`method` a string either `"KY"` for index ${}^{\alpha}K$ (if `disORtree` is of class `dist`) or ${}^{\alpha}Y$ (if `disORtree` is of class `phylo`, `phylo4`, or `hclust)`; or "KstarI" for index ${}^{\alpha}K^*$ (if `disORtree` is of class `dist`) or ${}^{\alpha}I$ (if `disORtree` is of class `phylo`, `phylo4`, or `hclust`). If several values are given, only the first one is considered.

`palpha` a nonnegative numeric or a vector with nonnegative value(s) for parameter $\alpha$.

`equivalent` a logical. If `TRUE`, the diversity values are calculated in terms of equivalent number of species (see Tables 1 and 2 in main text).

`option` a string either `"asymmetric"` or `"symmetric"`. The parameter `option` is only used if `disORtree` is of class `phylo`, `phylo4`, or `hclust`. If `option="symmetric"`, the distance between two tips on a tree is defined as half the sum of branch length on the smallest path that connects the two species; while if `option="asymmetric"` the distance between a tip *i* and a tip *j* on a tree is defined as the sum of branch lengths between tip *i* and its most recent ancestor with tip *j*. If the tree is ultrametric, the two options are equivalent.

`dmax` a nonnegative numeric indicating the maximum possible dissimilarity between two species. `dmax` must be higher than or equal to the maximum observed dissimilarity between two species. `dmax` must be higher than the longest distance from tip to root if a tree is used in `disORtree`.

`tol` a numeric tolerance threshold: values between - `tol` and `tol` are considered equal to zero.

`x` an object of class `FPdivparam` obtained with function `FPdivparam` .

`legend` a logical. If `TRUE` a legend is given with the colour, the type of line (etc.) used to define the diversity curve of each community.

`legendposi` a string that gives the position of the legend to be passed to function `legend` of the base of R.

`axisLABEL` a string to display on the main axis of the plot to designate what we are measuring. The default is `"FP-diversity"`.

`type` a string to be passed to the graphic argument `type` of functions `plot` and `lines` used to draw the diversity curve of each community.

`col` vector of colours to be passed to the graphic argument `col` of functions `plot` and `lines` to define the colour of the diversity curve of each community.

`lty` vector of type of line (plain, broken etc.) to be passed to the graphic argument `lty` of functions `plot` and `lines` used to draw the diversity curve of each community.

`pch` type of point (open circle, close circle, square etc.) to be passed to the graphic argument `pch` of functions `plot` and `lines` used to draw the diversity level of each community.

... other arguments can be added and passed to the functions `plot` and `lines` used to draw the graphic.

***Value:***
If only one value of `palpha` is given, function `FPdivparam` returns a vector with the phylogenetic diversity of each community.
If more than one value of `palpha` is given, a list of two objects is returned:
`palpha` (the vector of values for `palpha)`;
`div` (a data frame with the phylogenetic diversity of each community calculated for all values of `palpha`).

The function `plot.FPdivparam` returns a graphic.

**C.2. Abundance-based measures of species' rarity, functional or phylogenetic distinctiveness and functional or phylogenetic effective originality**

The function distinctAb calculates parametric indices of species' rarity and functional or phylogenetic distinctiveness and effective originality as in Tables 1 and 2 of the main text.

***Usage:***

```
distinctAb(comm, disORtree, method = c("Q", "KY", "KstarI"), palpha
= 2,
    option = c("asymmetric", "symmetric"), tol = 1e-10)
```

***Arguments:***

`comm` a data frame or a matrix typically with communities as rows, species as columns and abundance as entry. Species should be labelled as in object `disORtree`.

`disORtree` an object inheriting the class `dist`, giving the FP-dissimilarities between species, or inheriting the class `phylo` (see package ape), `phylo4` (see package `phylobase`), or `hclust`, where species are tips.

`method` a string either; "Q" for the quadratic entropy; `"KY"` for index ${}^{\alpha}K$ (if `disORtree` is of class `dist`) or ${}^{\alpha}Y$ (if `disORtree` is of class `phylo`, `phylo4`, or `hclust)`; or "KstarI" for index ${}^{\alpha}K*$ (if `disORtree` is of class `dist`) or ${}^{\alpha}I$ (if `disORtree` is of class `phylo`, `phylo4`, or `hclust`). If several values are given, only the first one is considered.

`palpha` a nonnegative value for parameter $\alpha$. `palpha` is ignored if `method = "Q"`.

`option` a string either `"asymmetric"` or `"symmetric"`. The parameter `option` is only used if `disORtree` is of class `phylo`, `phylo4`, or `hclust`. If `option="symmetric"`, the distance between two tips on a tree is defined as half the sum of branch length on the smallest path that connects the two species; while if `option="asymmetric"` the distance between a tip *i* and a tip *j* on a tree is defined as the sum of branch lengths between tip *i* and its most recent ancestor with tip *j*. If the tree is ultrametric, the two options are equivalent.

`tol` a numeric tolerance threshold: values between - `tol` and `tol` are considered equal to zero.

***Value:***

If `palpha` $\leq 1$, then, the function returns a list of four objects of class `data.frame` (with communities as rows and species as columns):

- `TotContr` provides for each species in each community the effective originality multiplied by the relative abundance;
- `EffOriPres` provides for each species present in each community the effective originality; NA for absent species;
- `DistinctPres` provides for each species present in each community the distinctiveness; NA for absent species;
- `Rarity` provides for each species in each community the rarity (maximum rarity for absent species).

Else, the function returns a list with the three objects of class `data.frame` presented above and two additional objects also of class `data.frame`:

- `EffOriAll` provides for each species its effective originality compared with the composition of each community; even absent species have a value considering they have zero abundance so maximum rarity and considering their functional dissimilarity or phylogenetic dissimilarity with all species present in each community;

- `DistinctAll` provides for each species its distinctiveness compared with the composition of each community; even absent species have a value considering they have zero abundance so maximum rarity and considering their functional dissimilarity or phylogenetic dissimilarity with all species present in each community.

##### C.3. Index $^{\alpha}D$eq* and related indices

The function `distinctDis` calculates five indices of species' distinctiveness.

***Usage:***

```
distinctDis(dis, method = c("Rb", "AV", "FV", "NN", "Dstar",
    "full"), palpha = 0, standardized = FALSE)
```

***Arguments:***

`dis` an object of class `dist` containing pair-wise (functional or phylogenetic) dissimilarities between species.

`method` a string or a vector of strings. Possible values are "Rb", "AV", "FV", "NN", "Dstar" and "full". "Rb" is for Pavoine et al. (2017) index Rb; "AV" is for AV, the average dissimilarity between a species and all others in a set (Eiswerth and Haney, 1992; Ricotta, 2004); "FV" is for FV, the average dissimilarity between a species and any other (including the focal species itself) (Schmera et al., 2009); "NN" is for the minimum dissimilarity between a species and any other (the dissimilarity to its Nearest Neighbor) (Pavoine et al., 2017); "Dstar" is the parametric indices $^{\alpha}D$eq* developed in the main text (see Appendix B); "full" returns all indices.

`palpha` a numeric or a numeric vector which provides the values of parameter $\alpha$ in index $^{\alpha}D$eq* developed in the main text (see Appendix B).

`standardized` a logical. If `TRUE`, the vector of originalities is divided by its sum (transforming absolute distinctiveness values into relative distinctiveness values).

***Value:***

A data frame with species as rows and distinctiveness indices as columns.

##### C.4. Index $^{\alpha}\Delta$eq* and related indices

The function `distinctTree` calculates indices of species' distinctiveness that rely on the structure and branch lengths of (phylogenetic) trees. Trees with polytomies are allowed.

***Usage:***

```
distinctTree(phyl, method = c("ED", "ES", "Delta*"), palpha = 0,
standardized = FALSE)
```

***Arguments:***

`phyl` an object inheriting the class `phylo` (see package `ape`), `phylo4` (see package `phylobase`) or `hclust`.

`method` a string or a vector of strings. Possible values are "ED", "ES", and "Delta*". "ED" is for the evolutionary distinctiveness, also named fair-proportion, index (Redding, 2003; Isaac et al., 2007); "ES" is for the Equal-Splits index (Redding and Mooers, 2006); "Delta*" is for index ${}^{\alpha}\Delta eq^{*}$ developed in the main text (see also Appendix B).

`palpha` a nonnegative value or a vector with nonnegative value(s) for parameter $\alpha$ of ${}^{\alpha}\Delta eq^{*}$.

`standardized` a logical. If `TRUE`, the vector of originalities is divided by its sum (transforming absolute distinctiveness values into relative distinctiveness values).

***Value:***
A data frame with species as rows and distinctiveness indices as columns.

##### C.5. Scripts used in the main text

```
install.packages("adephylo") # Jombart et al. (2010)
install.packages("adiv") # Pavoine et al. (2020)
install.packages("ape")
install.packages("phylobase")

library(adephylo)
library(adiv)
library(ape)
library(phylobase)

phy <- read.tree(file.choose()) # select Appendix D
ab <- read.table(file.choose(), header = TRUE, row.names = 1)[,
phy$tip.label] # select Appendix E

source(file.choose()) # select Appendix F

# Figure 3 of the main text:

divY <- FPdivparam(comm = ab, disORtree = phy, palpha=seq(0, 3,
length=100), equivalent = TRUE)
plot(divY, col=c(1,2,4, 5), pch=c(15:18))
title("Phylogenetic diversity, Index alphaY")
```

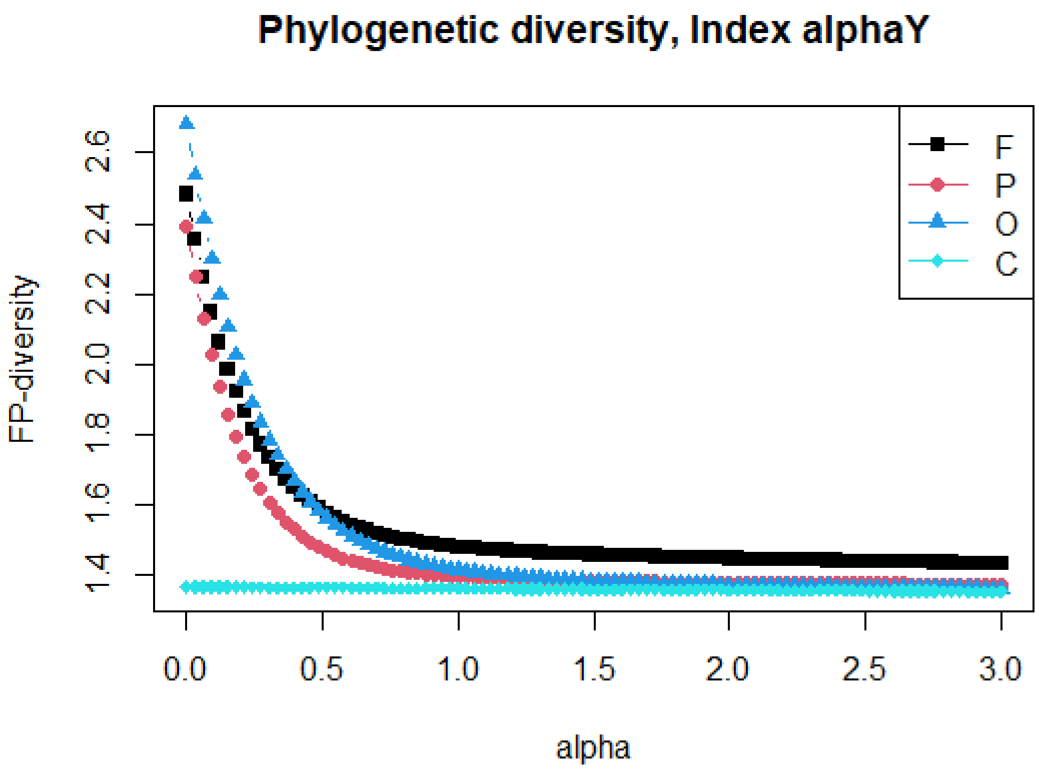


### F = rainforest
### P = cacao plantation
### O = old fields
### C = cornfields

```
divI <- FPdivparam(comm = ab, disORtree = phy, method = "KstarI",
palpha=seq(0, 3, length=100) , equivalent = TRUE)
plot(divI, col=c(1,2,4, 5), pch=c(15:18))
title("Phylogenetic diversity, Index alphaI")
```

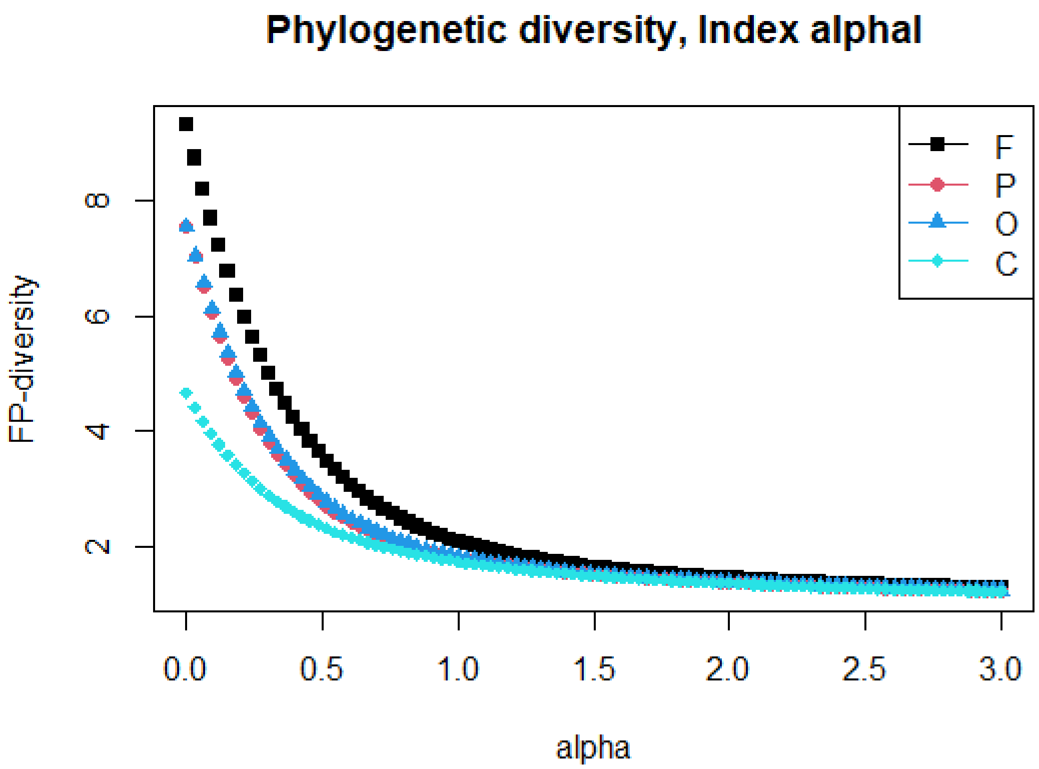


*# Figure 2 of the main text:*

```
U <- distinctTree(phy, method = c("ED", "Delta*"), palpha = c(-3, -
2, -1, 0, 1, 2, 3))
U.4d <- phylo4d(phy, as.matrix(U))
dotp4d(U.4d, center = FALSE, scale = FALSE, trait.labels = c("ED",
"-3Delta*", "-2Delta*", "-1Delta*", "0Delta*", "1Delta*", "2Delta*",
"3Delta*"))
```

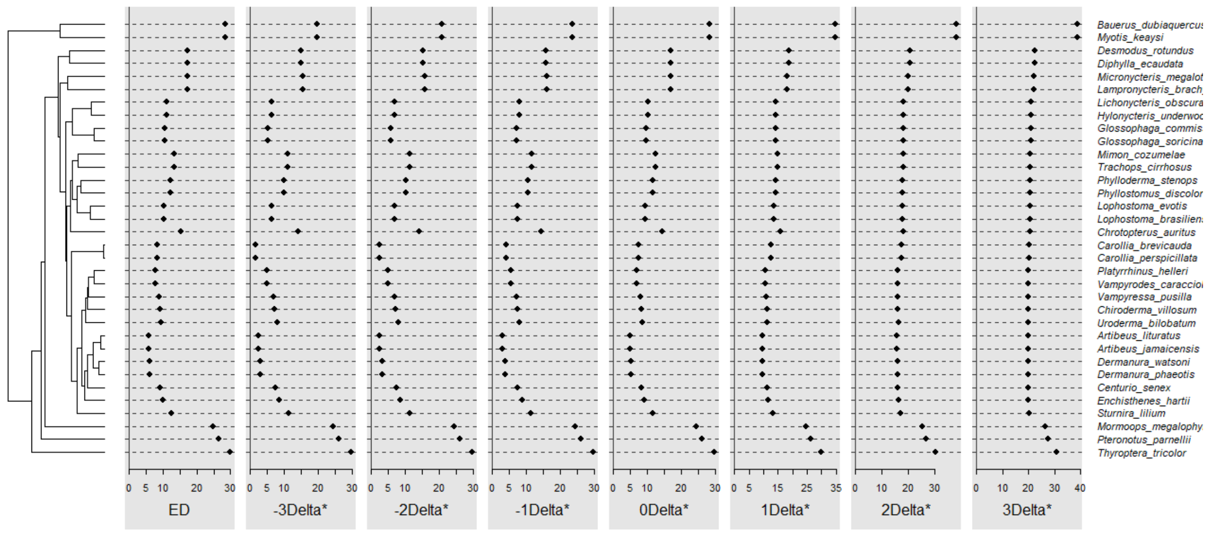

***# Figure 4 of the main text:***

```
REStotalQ <- distinctAb(ab, phy)
REStotalKstar0 <- distinctAb(ab, phy, method="KstarI", palpha=0)
REStotalKstar1 <- distinctAb(ab, phy, method="KstarI", palpha=1)

MATT <- as.matrix(cbind.data.frame(t(REStotalKstar0[[2]]),
t(REStotalKstar1[[2]]), t(REStotalQ[[2]])))
colnames(MATT) <- paste("v", 1:12)
U.4d <- phylo4d(phy,MATT)
dotp4d(U.4d, center = FALSE, scale = FALSE,
    data.xlim = matrix(c(rep(range(MATT[, 1:4], na.rm=TRUE), 4),
    rep(range(MATT[, 5:8], na.rm=TRUE), 4),
    rep(range(MATT[, 9:12], na.rm=TRUE), 4)), 2, 12),
    trait.labels = c("F, alpha=0", "P, alpha=0", "O, alpha=0",
    "C, alpha=0", "F, alpha=1", "P, alpha=1", "O, alpha=1",
    "C, alpha=1", "F, alpha=2", "P, alpha=2", "O, alpha=2",
    "C, alpha=2"))
title("Effective originality")
```

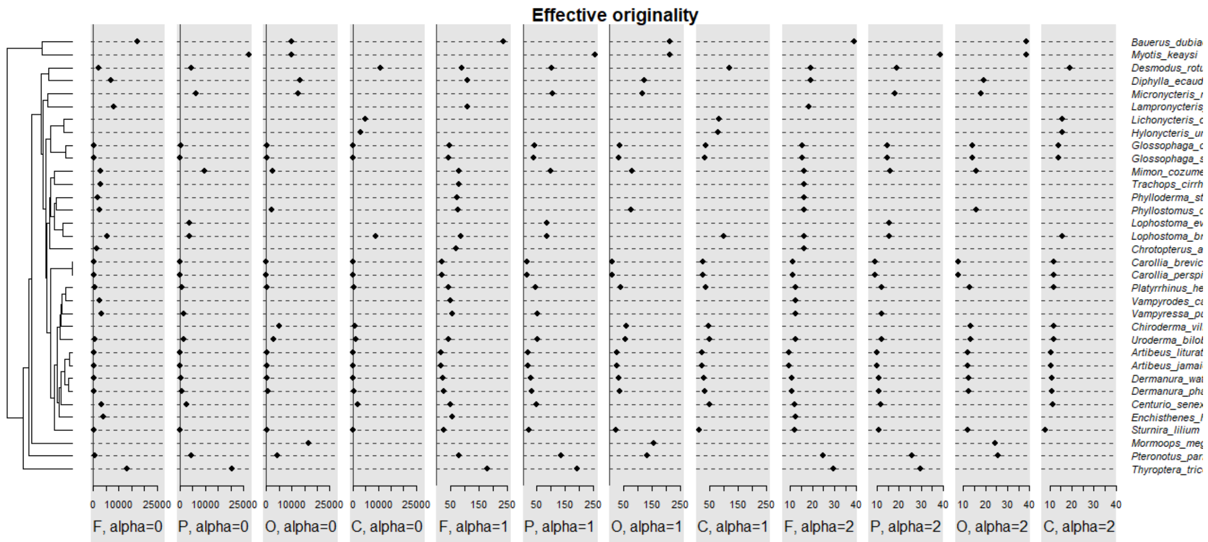


```
MATT2 <- as.matrix(cbind.data.frame(t(REStotalKstar0[[3]]),
t(REStotalKstar1[[3]]), t(REStotalQ[[4]])))
colnames(MATT2) <- paste("v", 1:12)
U.4d2 <- phylo4d(phy,MATT2)
dotp4d(U.4d2, dot.pch = 17, dot.col="blue", dot.cex=1.2,
    center = FALSE, scale = FALSE,
    data.xlim =   matrix(c(rep(range(MATT2[, 1:4], na.rm=TRUE), 4),
    rep(range(MATT2[, 5:8], na.rm=TRUE), 4),
    rep(range(MATT2[, 9:12], na.rm=TRUE), 4)), 2, 12),
    trait.labels = c("F, alpha=0", "P, alpha=0", "O, alpha=0",
    "C, alpha=0", "F, alpha=1", "P, alpha=1", "O, alpha=1",
    "C, alpha=1", "F, alpha=2", "P, alpha=2", "O, alpha=2",
    "C, alpha=2"))
title("Distinctiveness")
```

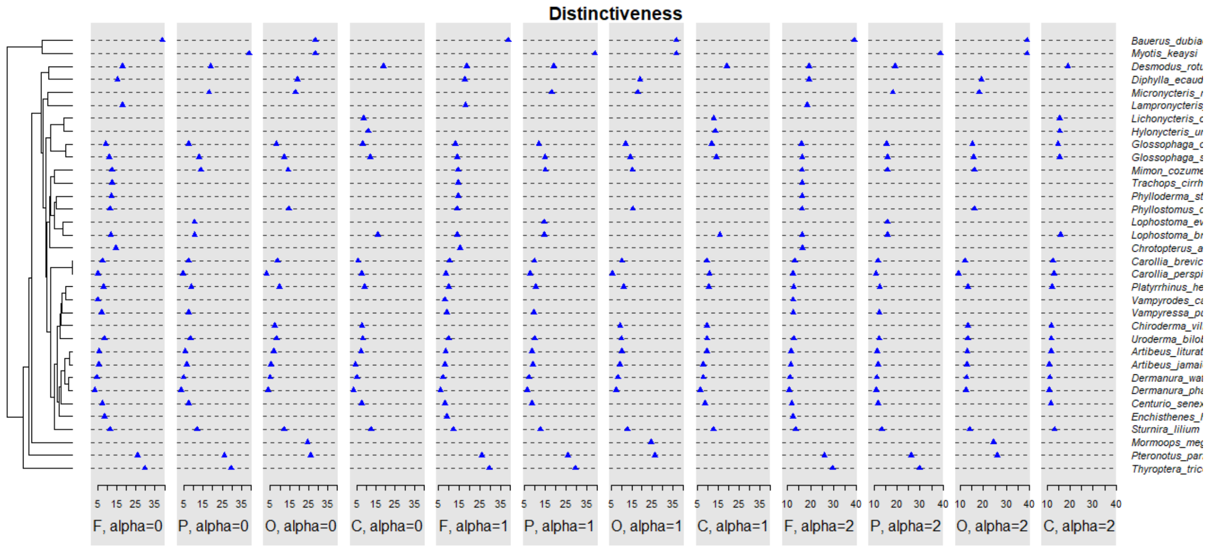
